# Flow molecular dynamics simulations reveal mechanosensitive regulation of von Willebrand factor through glycan-modulated autoinhibitory modules

**DOI:** 10.64898/2026.04.04.716521

**Authors:** Naveen E. Louis, Yunduo C. Zhao, Lining A. Ju

**Affiliations:** School of Biomedical Engineering, The University of Sydney, Darlington, NSW 2008, Australia; Charles Perkins Centre, The University of Sydney, Camperdown, NSW 2006, Australia; The University of Sydney Nano Institute (Sydney Nano), The University of Sydney, Camperdown, NSW 2006, Australia; Heart Research Institute, Camperdown, Newtown, NSW 2042, Australia

**Keywords:** Von Willebrand factor, Mechanobiology, Molecular dynamics simulation, Structural biology, Biophysics

## Abstract

Force-induced protein conformational changes govern many essential biological processes, yet their molecular mechanisms remain difficult to resolve. Von Willebrand factor (VWF), a central regulator of haemostasis, is activated by hydrodynamic forces in blood flow, but how mechanical signals propagate across its multidomain architecture is poorly understood. Here, we use flow molecular dynamics (FMD), a simulation framework that applies fluid forces via controlled solvent flow to interrogate mechanosensitive proteins. Using VWF as a model system, we reconstructed the complete mechanomodule (D′D3–A1–A2–A3; 1,109 residues) with native glycosylation by integrating crystallographic data and AlphaFold predictions. FMD simulations capture a force-driven transition from a compact, autoinhibited “bird’s-nest” ensemble to an extended, activated state, revealing asymmetric autoinhibitory strengths within the N′AIM and C′AIM modules of the A1 domain. By directly linking static structures to dynamic, force-regulated behaviour, this work establishes a generalizable platform for dissecting protein mechanosensitivity and enabling the rational design of force-responsive therapeutics.

**Graphical Abstract:** 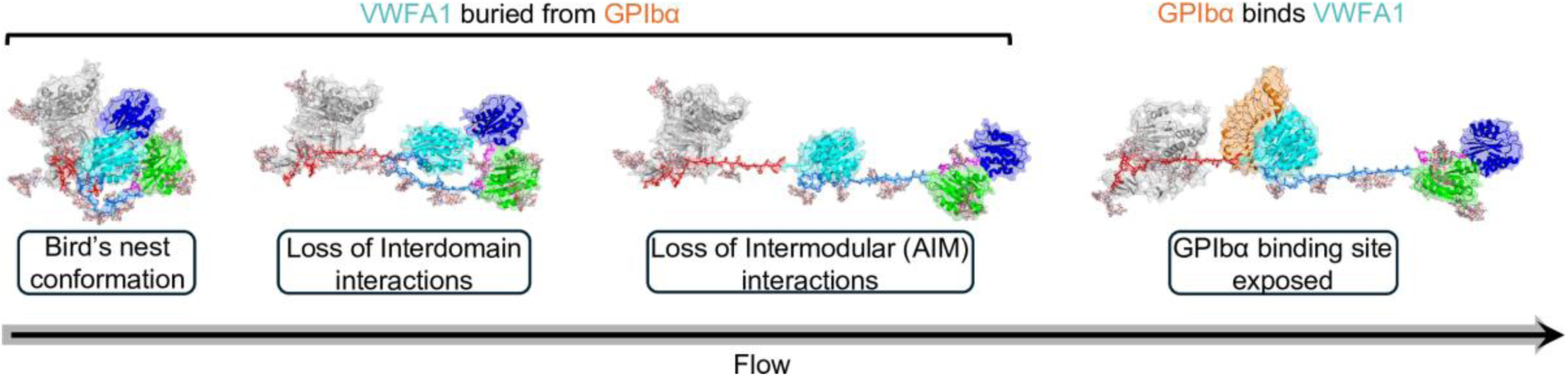

Flow molecular dynamics simulations reveal that GPIbα engages the A1 domain only after the disruption of key interdomain and intermodular interactions.

## Introduction

Von Willebrand Factor (VWF) exemplifies a finely tuned mechanosensitive protein system in which force-dependent conformational transitions directly govern its essential role in hemostasis^1,2^. Under physiological flow, VWF adopts a compact, globular “bird’s nest” conformation that conceals its platelet-binding domains, thereby preventing spontaneous adhesion^3,4^. However, at sites of vascular injury, elevated shear stress acts as a mechanical cue that triggers VWF to unfurl into an extended, conformation exposing cryptic binding sites for the platelet surface receptor glycoprotein Ibα (GPIbα)^5^. Remarkably, the spatial organization of VWF is highly context dependent. Within the trans-Golgi network, VWF monomers assemble via head-to-head interactions through the D’D3 domains and tail-to-tail associations via their C-terminal regions, forming higher-order multimers with a characteristic bouquet-like architecture^6^**(Fig. 1A, left)**. These multimers are compacted and stored within helical tubules inside Weibel–Palade bodies (WPBs), specialized endothelial granules that maintain VWF in a densely packed, secretion-ready state^7^ **(Fig. 1A, middle)**. Upon vascular injury, WPBs undergo rapid exocytosis, releasing VWF into the bloodstream, where these tubules unfurl into elongated multimers that tether platelets and initiate thrombus formation^8^ **(Fig. 1A, right)**. Although the functional consequences of this structural transition are well-established, the precise molecular architecture and mechanics governing VWF’s force-induced unfurling remain poorly understood representing a frontier in vascular mechanobiology.

**Figure 1.**
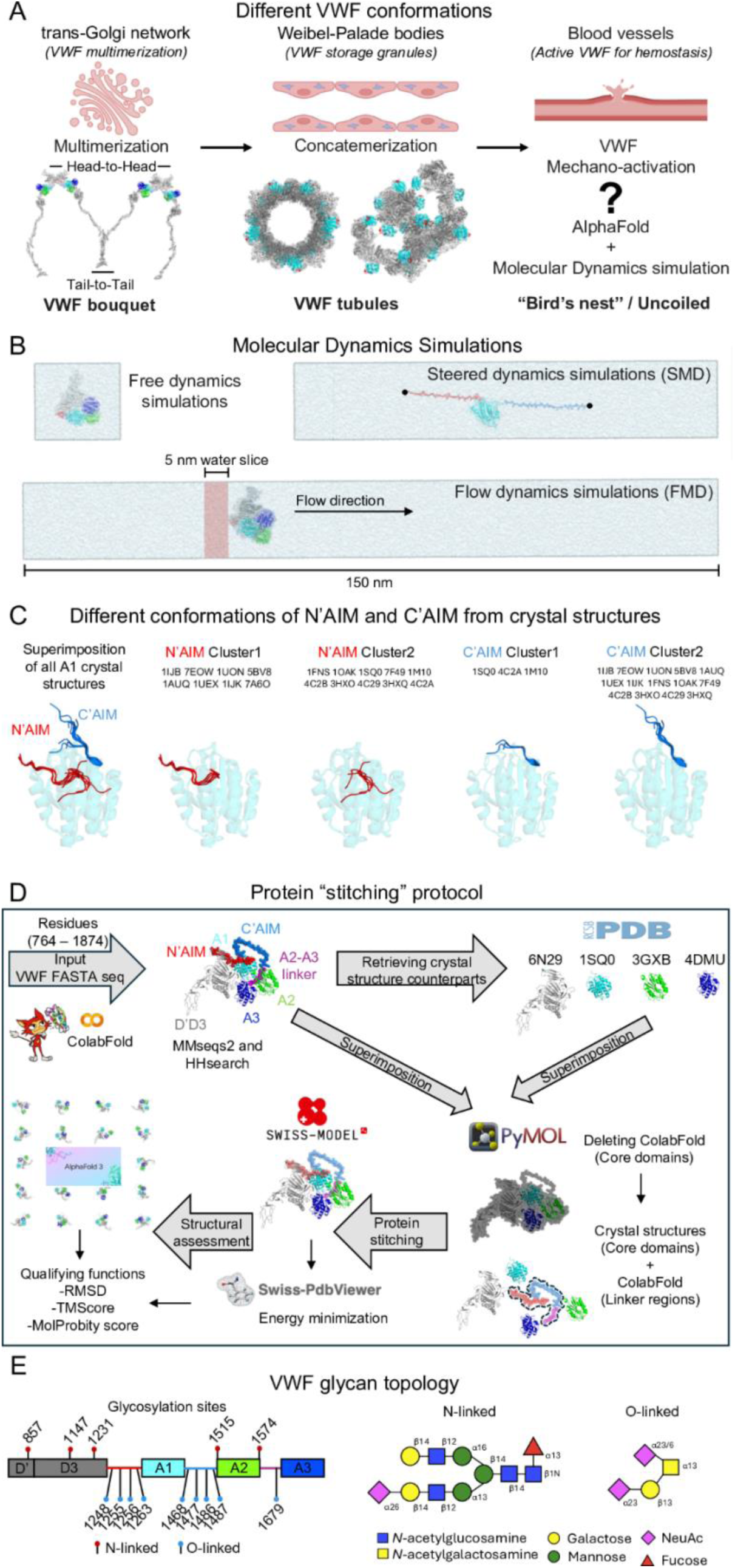
Conformational transitions underlying mechanoactivation of VWF and overview of the molecular modeling workflow and structural organization of the VWF mechanomodule. (A) VWF monomers assemble into ultra-large multimers within the trans-Golgi network through head-to-head dimerization mediated by the D’D3 domains and tail-to-tail interactions at their C-terminal cystine knot regions. These multimers are densely packed as helical tubules in Weibel–Palade bodies (WPBs) for intracellular storage. Upon vascular injury, WPBs undergo exocytosis, releasing VWF into the circulation, where the helical tubules unfurl under shear flow to expose platelet-binding sites, initiating platelet adhesion and thrombus formation. (B) Representative simulation strategies employed in this study, including equilibrium (free), flow-conditioned, and steered molecular dynamics, used to interrogate VWF mechanoactivation pathways. (C) Structural clustering of all available crystal structures of the A1 domain, illustrating conformational diversity in the N-terminal AIM (N’AIM) and C-terminal AIM (C’AIM) motifs. (D) Protein reconstruction strategy combining ColabFold-based predictions for unresolved linker regions with homology modeling guided by existing crystal structures. This integrative approach enables high-resolution atomic modeling of the VWF mechanomodule. (E) Left: Mapping of N- and O-linked glycosylation sites across the VWF sequence, highlighting their spatial distribution and occupancy. Right: Representative glycan topology used in simulations.

Recent breakthroughs in AI-guided structural biology, exemplified by AlphaFold 3, have revolutionized our ability to visualize complex protein architectures^9^. These tools provide unparalleled insights into static conformations and offer unprecedented opportunities to explore large, multidomain systems such as VWF. However, their static predictions limit their ability to capture force-induced transitions and dynamic rearrangements. To overcome these limitations, we subjected AI-derived models to a suite of molecular dynamics (MD) simulations **(Fig. 1B)**, enabling us to explore the full mechanical landscape of VWF from its compact, autoinhibited conformation to the force-activated state required for platelet adhesion.

The compact, autoinhibited conformation of VWF represents a state of functional stasis^6^, maintained through a hierarchical network of autoinhibitory mechanisms. The A1 domain central to platelet recruitment is subjected to two key levels of inhibition: (i) interdomain shielding by flanking domains such as D’D3 domain^10^ and A3 domain^11^ and (ii) intermodular autoinhibition by the N-terminal and C-terminal autoinhibitory modules (N’AIM, residues 1238–1271, and C’AIM, residues 1459–1493), which sterically occlude the GPIbα-binding interface^4^. However, the structural basis of this autoinhibition has remained elusive due to technical limitations in crystallography. Most AI-based prediction tools, such as AlphaFold, are trained on existing high-resolution crystal structures, and thus struggle to accurately model flexible or unresolved regions such as the N’AIM and C’AIM modules which are absent from all currently deposited A1 crystal structures **(Fig. 1C)**. We employed a hybrid strategy that combines deep learning-based protein structure prediction with homology modeling, yielding a more complete and functionally relevant representation of the autoinhibited VWF ensemble **(Fig. 1D)**. Furthermore, experimental efforts to produce AIM-A1 constructs with homogeneous O-glycosylation and resolve them at high resolution have been unsuccessful^12^. Given emerging evidence that glycosylation modulates VWF mechanosensitivity and spatial conformation^13,14^ **(Fig. 1E)**, our study incorporates N- and O-linked glycans associated with the VWF mechanomodule. This comprehensive modeling enabled us to explore how glycan-mediated shielding influences A1 accessibility. We also corroborate prior experimental evidence for the regulatory roles of N′AIM and C′AIM, while providing structural insight into why N′AIM exerts a more dominant autoinhibitory effect. Detailed steric-clash analyses mechanistically explain the differential binding affinities of distinct VWF constructs to GPIbα. Together, these findings refine our understanding of VWF mechanoregulation and establish a framework for mechanomedicine-guided drug design^15^ by delineating which VWF epitopes are shielded under static conditions and selectively exposed under force, i.e mechanosensitive mapping.

## Results

### Hybrid modeling and simulation of VWF reveals high confidence autoinhibited structure

The structural elucidation of glycoproteins such as VWF remains profoundly challenging, owing to the intrinsic chemical heterogeneity and dynamic conformational landscape imposed by their glycan moieties^16^. These complexities have significantly hindered efforts to resolve extended segments of VWF through crystallography. This limitation is reflected in the PDB, where nearly all deposited VWF structures comprise either isolated domains or discontinuous fragments, frequently omitting critical interdomain linker regions. Notably, the longest crystallized N’AIM in PDB entry 1U0N ^17^ only extends from Asp1260, thereby excluding the structurally unresolved but functionally essential distal segment spanning residues 1238–1259. Similarly, the most extended crystallised C’AIM captured to date PDB 5BV8 ^18^ terminates at Leu1469, omitting the distal residues 1470–1493 that are vital to understanding A1 domain regulation in its native context. Our integrative approach of modelling the hybridized model (CF+SM) consisting of crystallographic cores and ColabFold-modeled linkers spanning from the D’D3 – A3 domain and achieved better scores for metrics such as RMSD and TM-scores against crystal structures **(Table. S1)** and MolProbity scores following 500 ns simulations **(Table. S2).** Our rationale for focusing on the VWF mechanomodule arises from the observation that predictive models of the N’AIM–A1–C’AIM assembly, in the absence of flanking domains D’-D3, A2, and A3, yield linker regions in an artificially extended, linear conformation **(Fig. S1A).** Moreover, each structural prediction results in a single, static representation of the mechanomodule **(Fig. S1B)**, failing to capture the dynamic conformational landscape intrinsic to VWF. These limitations underscore the necessity of integrating molecular dynamics (MD) simulations with structural predictions to resolve the full range of conformational transitions exhibited by VWF, both in the absence and presence of external mechanical forces. We further validated our model by closely examining the N’AIM–A1–C’AIM region to assess whether the orientation of the flanking autoinhibitory modules aligns with known structural data **(Fig. S2A)**. Specifically, we compared our models against the A1 domain crystal structure (PDB ID: 1U0N), which contains the longest experimentally resolved N’AIM segment. The top-scoring AlphaFold 3 model and our CF+SM model both recapitulated similar poses for the structured portions of N’AIM (residues 1261–1271) and C’AIM (residues 1458–1468). However, notable variability persisted in the more flexible regions of N’AIM (residues 1238–1260) and C’AIM (residues 1469–1493). To determine which conformations may represent physiologically relevant autoinhibited states, we superimposed each model onto the A1–GPIbα complex structure (PDB ID: 1SQ0). This allowed us to evaluate the extent of steric hindrance imposed by N’AIM and C’AIM on the GPIbα-binding site. The top scoring AlphaFold 3 model **(Fig. S2B)** showed minimal obstruction of the GPIbα interface. In contrast, our CF+SM **(Fig. S2C)** and glycosylated CF+SM **(Fig. S2D)** models exhibited pronounced N’AIM-mediated shielding, consistent with the structural features expected of the compact, autoinhibited ‘bird’s nest’ conformation.

### Glycosylation modulates VWF mechanomodule assembly and dynamics

The presence of five N-linked and nine O-linked glycans led to distinct assembly patterns of the VWF mechanomodule **(Movie S3)** and increased the overall hydrodynamic size of VWF **(Fig. 2A).** To dissect the structural consequences of glycosylation, we performed molecular dynamics trajectory analyses for both glycosylated and non-glycosylated models during the initial (R_1_: 0–100 ns) and final (R_5_: 400–500 ns) stages of simulation **(Table S3)**. In the absence of glycans, the mechanomodule showed minimal structural deviation, with an RMSD of 0.26 nm at R_5_ compared to 0.38 nm in the glycosylated model, despite both averaging 0.97 nm during the R_1_ **(Fig. 2B)**. Glycosylation led to a less compact structural organization, with an average radius of gyration (Rg) of 3.75 nm, compared to 3.48 nm in the unglycosylated counterpart **(Fig. 2C)**. This reduced compactness was accompanied by greater solvent exposure, as reflected by a higher solvent-accessible surface area (SASA) of 575.95 nm² versus 523.45 nm² **(Fig. 2D)**. Notably, the number of intramolecular hydrogen bonds decreased in the glycosylated form (752 vs. 766), indicating weakened interdomain interactions and altered domain packing likely driven by glycan-induced steric hindrance. **(Fig. 2E)**. In addition, glycosylation increased the structural flexibility of the mechanomodule, with average Cα atomic fluctuations rising from 0.17 nm to 0.22 nm **(Fig. 2F)**. Detailed analysis of interdomain (D’-D3–A2–A3–A1) and intermodular (N’AIM–A1, C’AIM–A1, N’AIM–C’AIM) HB and SB revealed that glycosylation of the VWF mechanomodule attenuates interdomain interactions **(Fig. S3A)**, while preserving intermodular contacts (AIM–A1) **(Fig. 2G)**.

**Figure 2.**
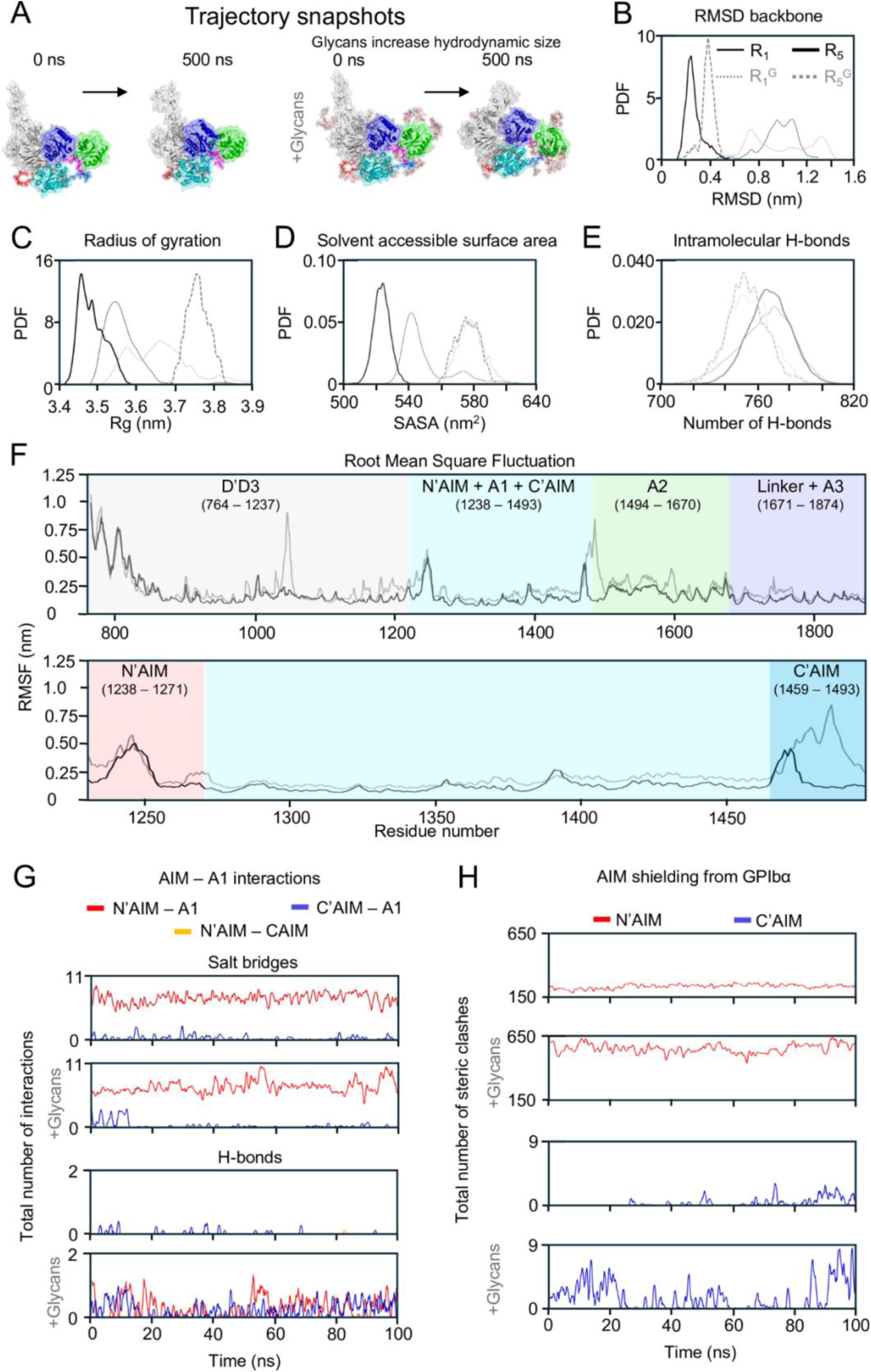
Free molecular dynamics simulations reveal the compact ‘bird’s nest’ conformation of VWF and the role of glycans in modulating structural dynamics. **(A)** Representative trajectory snapshots at t = 0 ns and t = 500 ns for glycosylated and non-glycosylated VWF mechanomodules, showing that glycans increase the overall hydrodynamic size. **(B–E)** Probability density distributions comparing glycosylated and non-glycosylated mechanomodules across structural metrics: **(B)** backbone root-mean-square deviation (RMSD), **(C)** radius of gyration (Rg), **(D)** solvent-accessible surface area (SASA), and **(E)** number of intramolecular hydrogen bonds. **(F)** Root-mean-square fluctuations (RMSF) of Cα atoms across the full mechanomodule (top) and focused analysis of the N’-AIM–A1–C’-AIM region (bottom), comparing late-stage trajectories (R_5_ vs. R_5_^G^). **(G)** Intermolecular interaction analysis between AIM and the A1 domain. **(H)** Steric clash analysis of A1 shielding by AIM, calculated from interatomic distances <3.0 Å to assess potential GPIbα-binding interference. Key: Thin black line: R_1_ (0–100 ns, no glycans); Thin grey dashed line: R_1_^G^ (0–100 ns, with glycans); Thick black line: R_5_ (400–500 ns, no glycans); Thick grey dashed line: R_5_^G^ (400–500 ns, with glycans).

### Electrostatic and steric mechanisms govern autoinhibition by N’AIM and C’AIM

We identified that Glu1260 from the distal N’AIM, along with Glu1264 and Glu1269 from the proximal N’AIM, critically modulate the interaction affinity between N’AIM and the A1 domain. Similarly, Glu1463 of the C’AIM governs its association with A1. Notably, these residues are all negatively charged and form salt bridges with a cluster of positively charged residues on A1, including Arg1274, Arg1306, Arg1308, Arg1312, Arg1315, Arg1334, Arg1336, Arg1341, and Arg1374 **(Fig. S3B)**. This charge-based interaction is particularly significant, as GPIbα–A1’s physiological binding partner is itself highly negatively charged and engages A1 via these same positively charged regions. These findings suggest that the autoinhibitory mechanisms of both N’AIM and C’AIM are charge-dependent, wherein key acidic residues in the AIMs transiently shield the GPIbα-binding site on A1 through electrostatic complementarity corroborating previous observations ^19^. We observed a correlation between the number of interactions formed between the AIM regions and A1, and their relative autoinhibitory strength. N’AIM, which formed substantially more salt bridges with A1 than C’AIM **(Fig. S3B)**, demonstrated a greater capacity to sterically occlude the GPIbα-binding site. Remarkably, the presence of eight O-linked glycans further amplified the autoinhibitory potential of both N’AIM and C’AIM **(Fig. 2H)**, as glycan-induced steric hindrance significantly enhanced their ability to obstruct A1–GPIbα engagement, characterized by the higher number of steric clashes.

Insights from our flow simulations **(Movie S4)**, which recapitulate the flow-induced unfurling of VWF and the uncoiling of N’AIM and C’AIM to expose the A1 domain **(Fig. 3A)**, revealed that while O-linked glycans enhance steric shielding of A1 from GPIbα, they also modulate the stability of AIM–A1 interactions. Specifically, glycan-induced steric hindrance shortened the lifetimes of both N’AIM–A1 and C’AIM–A1 interactions **(Fig. 3B)**, leading to earlier uncoiling events compared to the unglycosylated system **(Fig. 3C)**. Importantly, the key residues mediating these interactions were conserved regardless of glycosylation status **(Fig. 3D)**, indicating that the observed differences arise primarily from sterics ^20^. Analysis of steric clashes under increasing flow revealed a progressive reduction in steric obstruction over time, consistent with A1 mechanoactivation. Notably, while the steric contribution of C’AIM diminished entirely as unfolding progressed, N’AIM continued to impose steric hindrance, further supporting its dominant role in shielding A1 from GPIbα binding, an observation reported also in previous studies ^14^ **(Fig. 3E)**.

**Figure 3.**
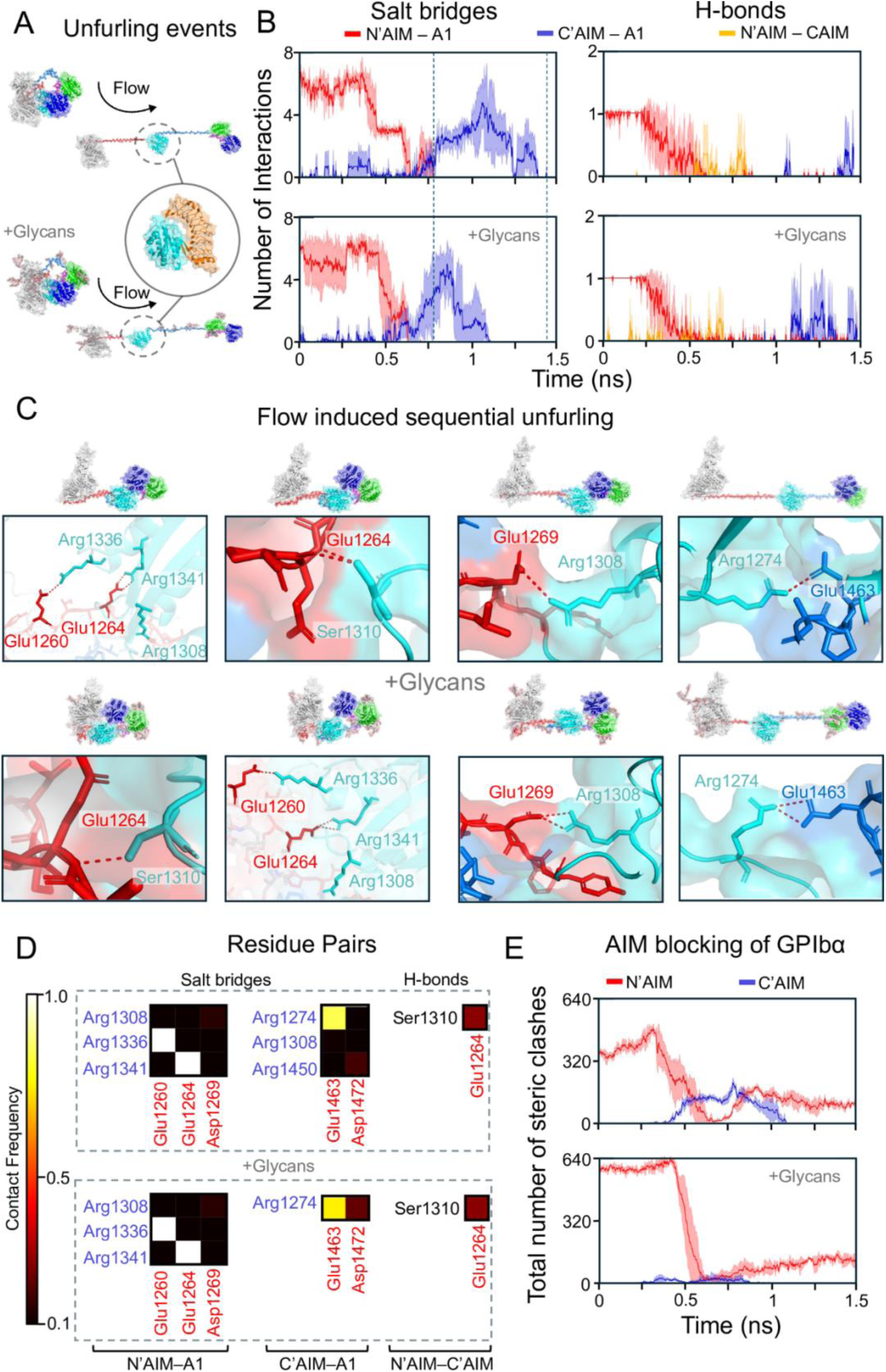
Flow-induced molecular dynamics (Flow-MD) simulations reveal the unfurling process of the VWF mechanomodule and its regulation of GPIbα accessibility. **(A)** Force-induced transition of the VWF mechanomodule from a compact “bird’s nest” conformation to an extended, uncoiled structure facilitates exposure of the A1 domain for GPIbα binding. **(B)** Dynamic intermolecular interactions between the A1 domain and the flanking N′AIM and C′AIM regions are illustrated, highlighting their regulatory role during unfolding. Time-resolved interaction profiles are shown as mean ± standard deviation, smoothed using the Savitzky–Golay filter. **(C)** Trajectory snapshots of key residues involved in modulating N’-AIM and C’-AIM engagement with A1, which together influence the shielding of the GPIbα-binding interface. **(D)** Contact frequency heatmap showing residue-residue interactions that persist in more than 10% of simulation frames; low-frequency contacts are excluded. Brighter colors indicate higher contact frequencies, while darker regions represent less persistent interactions. **(E)** Steric clash analysis of the N’-AIM and C’-AIM regions during unfolding, assessing their capacity to shield the A1 domain from GPIbα through atomic overlap (distance <3.0 Å) smoothed using the Savitzky–Golay filter.

To complement our flow MD simulations which mimic shear forces through solvent-mediated mechanical perturbation, we also conducted steered MD simulations **(Movie S5)**, applying tensile forces directly through the VWF backbone to assess differences in AIM-mediated shielding of A1 **(Fig. S1C)**. As shown in **Fig. 4A**, glycan-induced steric bulk reduced the spatial proximity of both N’AIM and C’AIM to A1, yet these sterics remained effective in obstructing GPIbα access **(Fig. 4B)**. Notably, the mean number of steric clashes contributed by C’AIM to GPIbα blocking increased from 0 in the non-glycosylated state to 1.54 with glycans. However, this effect is force-regime dependent: in Steered MD, the applied tension pulls C’AIM away from N’AIM, reducing its shielding efficacy. In contrast, flow MD promotes a more distributed uncoiling of N’AIM and C’AIM without backbone-directed tension, allowing C’AIM to maintain closer proximity to A1 for longer durations. This highlights the differential mechanical outcomes imposed by distinct force application paradigms and underscores the context-specific role of AIM in regulating A1 accessibility.

**Figure 4.**
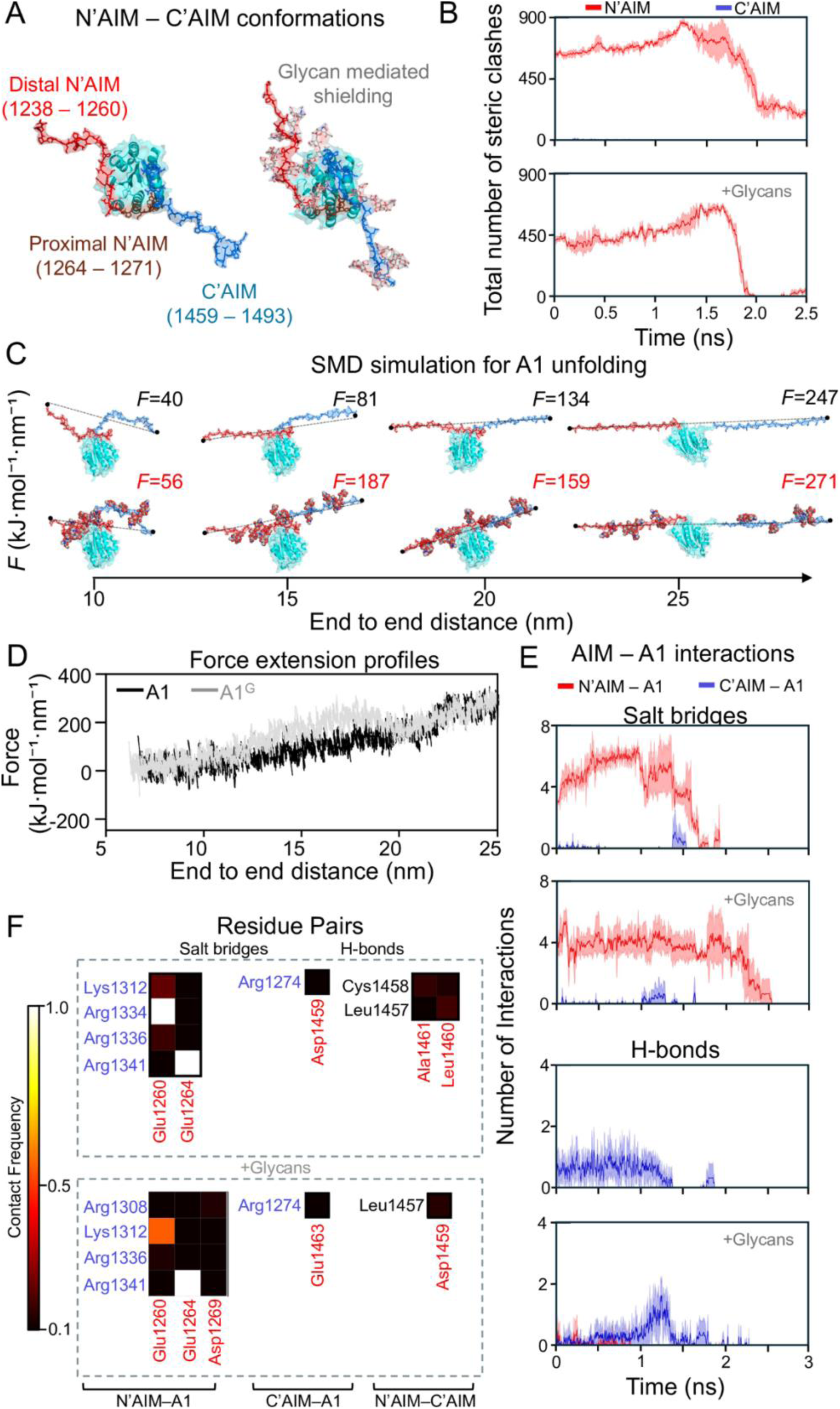
Steered molecular dynamics (Steered-MD) simulations reveal the force-induced uncoiling of the N’-AIM and C’-AIM regions and their role in A1 shielding under tensile stress. **(A)** Structural differences in the conformations of N’-AIM and C’-AIM with and without glycans, highlighting the spatial separation between their distal and proximal termini relative to the A1 domain. **(B)** Steric clash analysis of the N’-AIM and C’-AIM regions during forced uncoiling, assessing their potential to shield the A1 domain from GPIbα via atomic overlaps (distance <3.0 Å), smoothed using the Savitzky–Golay filter. **(C)** Representative trajectory snapshots illustrating the force required to uncoil the mechanomodule at increasing distances between N’-AIM and C’-AIM, in both glycosylated and non-glycosylated contexts. **(D)** Force-extension profiles showing the mechanical resistance associated with uncoiling of the AIM regions under steered MD (A1 – without glycans; A1^G^ with glycans). **(E)** Intermolecular interactions between the AIM regions and the A1 domain during force application, revealing transient or persistent binding contacts that may regulate GPIbα accessibility, smoothed using the Savitzky–Golay filter. **(F)** Contact frequency heatmap showing residue-residue interactions that persist in more than 10% of simulation frames; low-frequency contacts are excluded. Brighter colors indicate higher contact frequencies, while darker colors denote weaker or transient interactions.

Applying tensile forces to pull the C′AIM away from the N′AIM revealed that a higher force is required in the presence of eight O-linked glycans, which likely contribute hydrodynamic drag during extension **(Fig. 4C)**. This results in an increased force threshold needed to achieve equivalent extension distances **(Fig. 4D)**. Despite fewer intermolecular interactions compared to unglycosylated systems **(Fig. 4E)**, these contacts persisted for longer durations in glycosylated constructs **(Fig. S4)**, with contact frequencies sustained over extended timescales **(Fig. 4E)**. Notably, interactions between N′AIM–A1 and C′AIM–A1 identified in flow simulations were similarly observed in steered MD simulations **(Fig. 3D**, **Fig. 4F)**, reinforcing the critical role of AIM–A1 affinity, which is predominantly governed by electrostatic forces.

MSA of VWF residues 1238–1493 across *Homo sapiens*, *Mus musculus*, and *Bos taurus* **(Fig. S5A)** revealed that the proximal N’AIM is significantly more conserved than the distal N’AIM, highlighting its evolutionary and physiological importance. Notably, the distal N’AIM showed no sequence consensus in *Bos taurus*, further emphasizing its species-specific variability. Among all identified glycosylation sites, only the O-linked glycans at Ser1263 and Thr1487 were not conserved. Integrating these sequence insights with our flow and steered MD simulations, we observed that proximal N’AIM acts as the final barrier, maintaining steric hindrance within 3.0 Å of GPIbα. Furthermore, the glycan at Ser1263 emerged as the last remaining glycan obstructing A1 from GPIbα engagement, reinforcing its functional importance in autoinhibition **(Fig. S5B)**.

### Mechanical Unmasking of GPIbα Binding Sites by the VWF D′D3 Region

To further corroborate the unfolding kinetics of VWF observed in our simulations, we leveraged previously published kinetic data from Madabhushi *et al.*^21^ examining multiple VWF constructs, including ΔPro-VWF, ΔD′D3-VWF, ΔD′D3NFP⁻-VWF, and ΔD′D3OG⁻-VWF **(Fig. 5A)**. Although these studies employed dimeric VWF constructs, we established correspondence between experiment and simulation by comparing the reported dissociation constants (K_d_) with steric-clash and accessibility analyses of GPIbα binding during force-induced unfolding in our flow-based molecular dynamics simulations **(Fig. 5B)**. Madabhushi *et al.* demonstrated that deletion of the D′D3 domain together with the N-terminal flanking peptide up to residue 1267 resulted in the highest binding affinity for GPIbα^21^ **(Fig. 5C)**, highlighting the critical shielding role of these regions. Consistent with these observations, our simulations reproduced the same affinity trend, revealing increased GPIbα accessibility upon force-mediated unmasking of these domains **(Fig. 5D)**. Collectively, our results demonstrate that flow-induced simulations can identify force-sensitive epitopes and delineate mechanically protected versus exposed regions of VWF, providing a mechanistic framework to guide the rational design of next-generation antithrombotic therapeutics.

**Figure 5.**
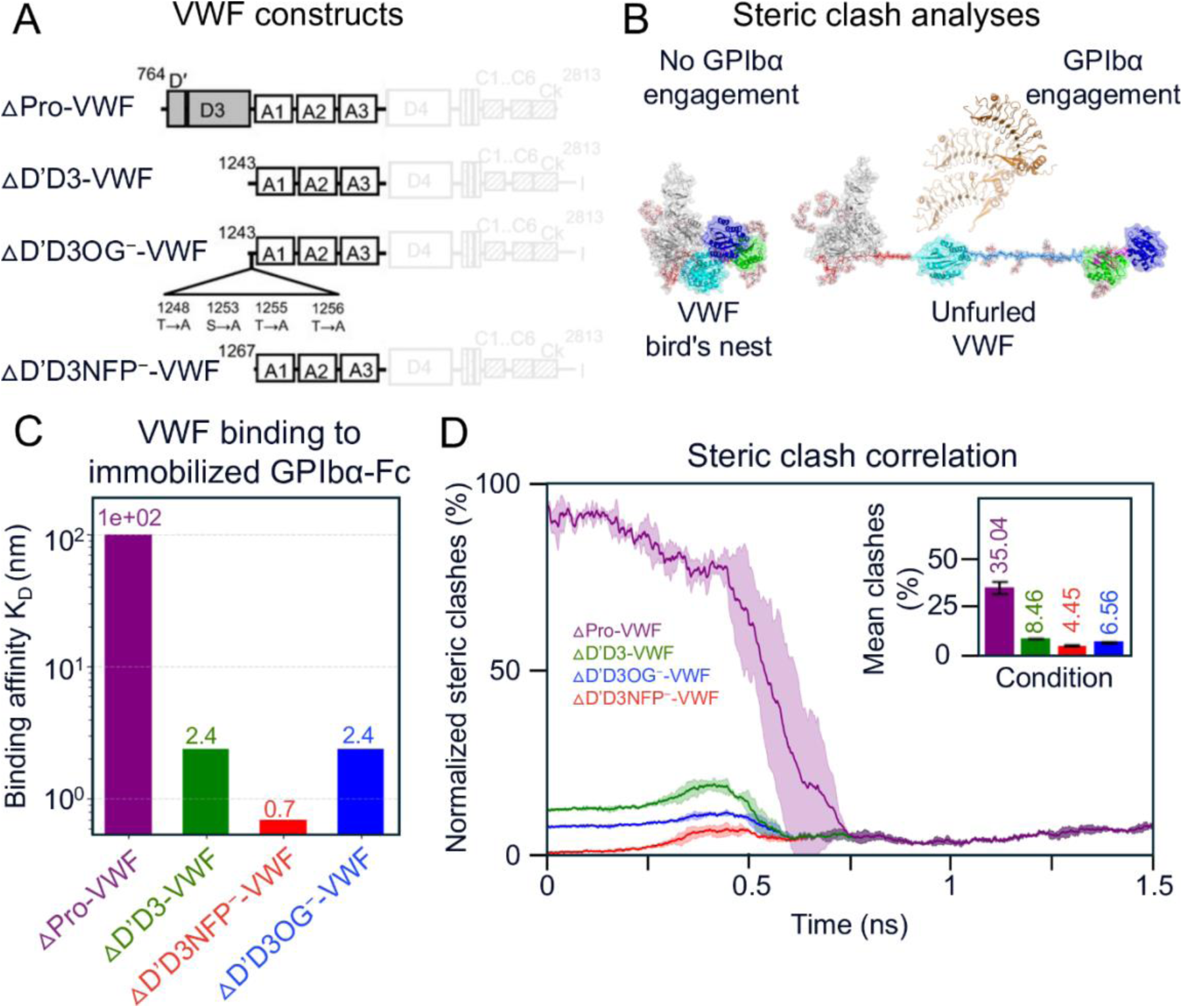
Kinetic correlation landscape of VWF–GPIbα affinity. (A) VWF constructs examined in Madabhushi et al^21^(B) Schematic of the steric-clash analysis used to evaluate GPIbα binding to VWF during force-induced unfolding.(C) Experimentally measured binding affinities reported by Madabhushi et al^21^(D) Normalized steric clash between GPIbα and distinct VWF regions during unfolding, smoothed using a Savitzky–Golay filter. The inset shows the mean steric-clash values.

## Discussion

VWF undergoes dramatic conformational transformations, transitioning from a compact ‘bird’s nest’ architecture to an elongated, uncoiled state in response to shear flow. Although neural network–based models have revolutionized the prediction of static protein structures, they remain inherently limited in capturing the dynamic flexibility and structural heterogeneity that define mechanosensitive proteins such as VWF ^22^. To transcend these limitations, MD simulations have become indispensable, offering atomistic resolution into VWF’s behaviour under tensile and flow conditions ^23–27^. These simulations yield mechanistic insights into VWF mchanomodulation, illuminating how mechanical forces modulate structure and, ultimately, biological function ^22^.We selected to simulate the D’-D3–A3 mechanomodule due to prior use in kinetic studies where spatial segregation of A1 domains was critical. Importantly, this construct retains the O-linked glycans and structural motifs proposed to regulate the accessibility and functional presentation of the A1 domain ^28^. Our free, flow-based, and steered MD simulations not only substantiated previous experimental findings but also revealed previously unrecognized mechanisms of VWF mechanomodulation, including dynamic interactions between the N′AIM and C′AIM regions and the A1 domain.

It is well established that glycans enhance the shielding of the A1 domain from GPIbα. Our steric clash analyses not only recapitulated this phenomenon but further demonstrated that the N′AIM region alone is sufficient to inhibit A1–GPIbα binding **(Figs. 2H, 3E, 4B, Table S4)** corroborating previous studies ^13,14^. Furthermore, we demonstrate that the interactions between N′AIM and the A1 domain is greater than that of C′AIM–A1**(Figs. 2G, 3D, 4F, Tables S5,S6)**, corroborating earlier findings^4^. This enhanced affinity is primarily driven by electrostatic interactions, including the formation of salt bridges, which contribute to the superior autoinhibitory capacity of N′AIM. Our steered MD simulations also corroborate previous findings that glycosylated forms of VWF exhibit greater mechanical stability and require higher forces to induce unfolding ^29^.

Interestingly, our simulations did not capture direct interactions between the distal regions of N’AIM and C’AIM, as previously proposed ^12^. This discrepancy likely stems from a key difference in simulation setup: we modeled A1 in the context of its flanking domains, D’D3 and A2, which impose spatial constraints that may hinder distal N’AIM–C’AIM interactions. In contrast, prior studies that simulated isolated N’AIM–A1–C’AIM constructs without neighbouring domains have reported contacts between the distal ends of these autoinhibitory modules ^30^. These observations suggest that the conformational pose and interaction landscape of N’AIM and C’AIM are strongly influenced by their structural context, and that domain architecture plays a critical role in regulating A1 autoinhibition ^31^.

Our simulations, complemented by MSA, reveal key regions within the N′AIM and critical glycans that act as the final steric barriers to GPIbα engagement during VWF extension. Prior studies have suggested that interactions between the proximal N′AIM and the A1 domain is essential for maintaining A1 in an autoinhibited conformation, disruption of these interactions is thought to activate A1, priming it for GPIbα binding ^12^. Our simulations identify specific residues most notably Glu1264 and Glu1269 within proximal N′AIM, that regulate N′AIM–A1 interactions **(Figs. 3D, 4F, S3B)**. In addition, Glu1260 from distal N′AIM was also observed to form contacts with A1 ^12^. MSA further revealed that the O-linked glycosylation site at Ser1263 is not conserved across species. In contrast, the proximal region of N’AIM, but not the distal segment exhibited relatively high sequence conservation, suggesting a potential role of these regions serving as the final shield of A1 from premature GPIbα engagement **(Fig. S5)**.

These findings underscore the utility of our analyses in elucidating VWF dynamics under both static conditions and flow-induced activation. Our approach provides atomic-level resolution to identify epitopes that are shielded in the quiescent state and selectively exposed under force. This paradigm is critical for rational drug design, enabling the prioritization of force-sensitive epitopes over those accessible in static structures, thereby facilitating efficient preclinical assessment prior to microfluidic experimentation and downstream validation.

## Methods

### *In silico* modeling of the VWF mechanomodule

To model the VWF mechanomodule in a physiologically relevant state and interrogate flow-induced activation, particularly the autoinhibitory roles of the N’AIM and C’AIM inhibitory modules of A1, we constructed an integrative structural model spanning residues 764–1872 (D′D3–A3 domains).Initial structural predictions were generated using AlphaFold3 ^9^, yielding a model with moderate interdomain confidence (predicted TM-score ≈ 0.482). To improve structural accuracy, we developed a hybrid modeling pipeline combining deep-learning-based predictions with high-resolution crystallographic templates. Linker regions absent in crystallographic **(Fig. 1C)** were reconstructed using ColabFold (CF), an accelerated AlphaFold2 implementation with MMseqs2 ^32,33^ for multiple sequence alignment. Experimentally resolved structures for D′D3 (PDB: 6N29), A1 (PDB: 1SQ0), A2 (PDB: 3GXB), and A3 (PDB: 4DMU) ^34–37^ were aligned to the CF scaffold. CF-predicted domains were replaced with their crystal counterparts to maintain empirical accuracy **(Fig. 2D)**, as AlphaFold predictions, although reliable, cannot supersede experimentally validated coordinates ^38^.The hybrid CF+SM model was assembled using PyMOL ^39^ and SWISS-MODEL (SM) ^40^ using the user template option, followed by energy minimization in Swiss-PdbViewer ^41^ to remove steric clashes and optimize stereochemistry. Model validation against AlphaFold3 and experimental templates was performed using TM-score, RMSD, and MolProbity ^42^ metrics.VWF contains five N-linked and nine O-linked glycosylation sites **(Fig. 1E, left)**. Structural models of glycans primarily disialylated core-1 O-glycans (∼70%) and monosialylated bi-antennary N-glycans (∼80%) ^43^ **(Fig. 1E, right)** were generated using the Glycan Reader & Modeler in CHARMM-GUI ^44^ and incorporated into the final atomistic model.

### Molecular dynamics simulation of VWF mechanomodulation: from compact to extended states

Equilibrium (Free) MD simulations – the “bird’s nest” conformation All molecular dynamics (MD) simulations were performed using GROMACS 2021.2 ^45^ with the CHARMM27 force field and TIP3P water model. All systems were neutralised by adding counter Na+ and Cl- ions using the genion tool. Systems were energy-minimized with the steepest descent algorithm and equilibrated in the NVT and NPT ensembles (100 ps each) at 300 K, using a 2 fs time step and the leap-frog integrator. Production simulations of 100 ns employed particle-mesh Ewald (PME) for long-range electrostatics and optimized nonbonded cutoffs following Lemkul et al. ^46^. Simulations with diverging backbone RMSD profiles were iteratively refined by restarting from the final frame until convergence was achieved. The same procedure was applied to glycosylated models.

Flow MD simulations – solvent-driven unfolding of the mechanomodule

To investigate the flow-induced unfurling and uncoiling of VWF, flow simulations were initiated from the stabilized conformation obtained at 500 ns of the free dynamics simulations. Earlier studies have introduced molecular dynamics frameworks to emulate flow-like environments, providing valuable insights into protein conformational transitions and informing the design of our approach ^23,24,47^. These studies also identified key methodological considerations particularly, that increasing flow velocity over time can restrict simulation duration and result in significant system heating, with temperatures reaching up to ∼350 K. Such thermal effects may compromise protein thermostability and secondary structure, potentially confounding interpretations of unfolding pathways. Notably, most earlier simulations examined short polypeptide segments typically ≤40 residues such as flexible loops or linker regions. While these systems are ideal for method development, they do not capture the structural complexity or mechanical resistance of large, multidomain proteins. In contrast, our study focused on a 1108-residue construct of VWF that includes well-defined α-helices and β-sheets, necessitating the application of a stronger but thermally stable flow field to ensure mechanical unfolding without introducing thermal artifacts. To investigate the effects of flow-like force application on VWF uncoiling, we explored solvent-based mechanical perturbation rather than applying tensile forces through steered MD simulations. Initially, we applied uniform acceleration to the oxygen atoms of water molecules at 10 nm ps⁻². Although this induced collective solvent drift, it failed to unfurl the protein **(Movie S1, top)**. In contrast, pulling water molecules toward the protein using an umbrella sampling bias effectively unfolded the mechanomodule **(Movie S1, bottom)**. This contrast stems from differences in force transmission: acceleration produces bulk water flow which, under periodic boundary conditions, exerts little net force on the protein due to reorganization and flow around the macromolecule. Umbrella pulling, by comparison, imposes a localized, directional force that persistently drives water into the protein surface, mimicking forced hydration and generating high-pressure contacts capable of disrupting hydration shells and intramolecular interactions. To isolate solvent-induced effects, the Cα atoms of the D′D3 domain were restrained with a harmonic potential (1000 kJ mol⁻¹ nm⁻²) in all simulations, preventing translational drift. Only the oxygen atoms of water molecules were pulled relative to the restrained domain to reduce high-frequency noise associated with lighter hydrogen atoms ^23^, allowing solvent-mediated shear to drive uncoiling of the VWFA1A2A3 domains without direct tensile stress on the backbone. The direction of flow was adapted from previous studies ^5^. The compact “bird’s nest” conformation of VWF was placed in an extended simulation box (15 × 15 × 150 nm) to allow elongation along the Z-axis. The system was solvated, neutralized, and equilibrated under standard conditions. Flow was applied by pulling a 5-nm slab of water oxygen atoms along the Z-axis using a harmonic spring potential (1000 kJ mol⁻¹ nm⁻²) at a constant velocity of ∼0.01 nm ps⁻¹. The applied force exceeds physiological shear rate values, but such accelerated conditions are standard in molecular dynamics to observe unfolding events within accessible simulation timescales This setup enabled controlled, stepwise uncoiling of VWF and characterization of its mechanotransductive response under flow **(Movie S2)**. Simulations were conducted in triplicates for the mechanomodule both with and without glycans to investigate unfolding patterns, the influence of interdomain interactions, and the role of AIM in exposing A1.

Steered MD simulations –glycan-mediated mechanical resilience

Steered MD (SMD) simulations were performed to probe glycan-mediated stabilization of A1, following established protocols ^27^. Each system was embedded in a 10 × 200 × 10 nm box with CHARMM36 protein/carbohydrate parameters and TIP3P water using CHARMM-GUI ^44^. A harmonic spring (1000 kJ·mol⁻¹·nm⁻²) was applied between the Cα atoms of Asn1493 and Gln1238, pulling along the Y-axis at 0.01 nm·ps⁻¹ to mimic atomic force microscopy or optical tweezer setups ^48^. Force–extension profiles were recorded, and rupture forces were defined as the peak force prior to detachment. Triplicate simulations were averaged to quantify glycan effects.

## Data analysis and visualization

Structural and dynamical analyses were conducted using GROMACS tools: gmx rms (RMSD) and gmx rmsf (root mean square fluctuation), gmx gyrate (radius of gyration), gmx sasa (solvent accessibility), and gmx hbond (hydrogen bonding) ^49,50^.Trajectories were sorted using MDTraj ^51^ and visualised in PyMOL, VMD, and QtGrace ^39,52,53^. The structural quality of the AlphaFold 3 model, the ColabFold–SWISS-MODEL hybrid (CF+SM), and the 500-ns simulation-refined CF+SM structure was assessed using MolProbity, providing comprehensive all-atom validation through integrated metrics including clashscore, rotamer outliers, and Ramachandran statistics ^54^.

Intermolecular interactions and steric clashes were analyses using MDAnalysis v 2.0 ^55^. Hydrogen bond (HB) and salt bridge (SB) interactions were evaluated from the MD trajectories. For H-bonds, donor atoms were defined as nitrogen or oxygen atoms capable of forming hydrogen bonds, specifically: N, NE, ND1, NE2, NH1, NH2, OG, OG1, and OH; acceptor atoms were defined as O, OD1, OD2, OE1, OE2, OG, OG1, and OH. A HB was considered present when the donor–acceptor distance was ≤ 3.5 Å and the angle between donor and acceptor atoms (approximating the donor–hydrogen–acceptor angle) was ≥ 150°, following established geometric guidelines. SB were defined as interactions between oppositely charged side chains, specifically between the positively charged nitrogen atoms of Lys (NZ) and Arg (NE, NH1, NH2) and the negatively charged oxygen atoms of Asp (OD1, OD2) and Glu (OE1, OE2). A SB was considered present when the distance between any such atom pair was ≤ 3.5 Å. Analyses were performed over every frame of the simulation trajectory, and interaction frequencies were recorded for quantitative comparison. Intermolecular interactions were analyzed by combining data from three independent replicate simulations. Each replicate’s trajectory frames were reindexed and concatenated to form a continuous dataset, allowing comprehensive assessment across all simulations. Contact frequencies were calculated as the fraction of frames in which each residue pair was observed to form a SB or HB across the concatenated dataset. To focus on persistent interactions, only contacts present in at least 10% of the combined frames were retained for further analysis and visualization as a heatmap.

MD trajectories were analysed against a static reference structure containing the GPIbα domain (PDB ID: 1SQ0) to quantify potential steric clashes. The static A1 domain from the reference structure was aligned to each frame of the trajectory using the heavy atoms of residues 1290–1430 as the alignment region. Following alignment, interatomic distances were computed between heavy atoms of the static GPIbα domain (residues 1–265) and dynamic regions of interest in the trajectory: NAIM (residues 1238–1271), CAIM (residues 1459–1493), including any N- or O-linked glycans within 5 Å of each region adapted from previous studies ^14^. Steric clashes were defined as any heavy atom pairs within 3.0 Å between the static reference and the dynamic trajectory. Clash counts were computed on a per-frame basis to enable quantitative assessment of glycosylation-dependent interference with GPIbα binding. Multiple sequence alignments (MSA) of the A1 domain from *Homo sapiens (NCBI Reference: AAB34053.1)*, *Mus musculus (NCBI Reference: ABC86574.1)*, and *Bos taurus (NCBI Reference:* NP_*001192237.1)* was performed, and residue conservation was analysed using Jalview ^56^.

## Limitations

Although our simulations provide a novel approach to understanding VWF unfolding kinetics, with corroboration from prior experimental studies, they are inherently limited by the scope of existing experimental data. Future studies examining the effects of VWF glycosylation and its differential interactions with therapeutic agents will be essential to validate and complement our in *silico* observations with experimental benchmarks.

## Supporting information

This PDF file includes Figures S1 to S5 and Tables S1 to S6

## Acknowledgments

The authors would like to thank Dr. Haimei Zhao and NCRI infrastructure and USYD SHM core research facility for computing resources installation. We thank Profs. Heyu Ni and Freda Passam for their clinical guidance and support. We thank Dr. Alex Depuy, Jianfang Ren and Catherine Chen for providing critical suggestions. Generative artificial intelligence tools were used solely for editorial assistance, including grammar and stylistic improvements. No AI tools were used for data generation, model development, statistical analysis, or scientific decision-making. All results and interpretations were produced and verified by the authors.

## Author Contributions

Designed research: L.A.J, N.E.L, Y.C.Z

Performed research: N.E.L

Analyzed data: N.E.L

Wrote the paper: L.A.J, N.E.L, Y.C.Z

## Competing Interest Statement

The authors declare no competing interests.

## Data availability

All data are available in the main text or the supplementary materials.

---

**Fig. S1.**
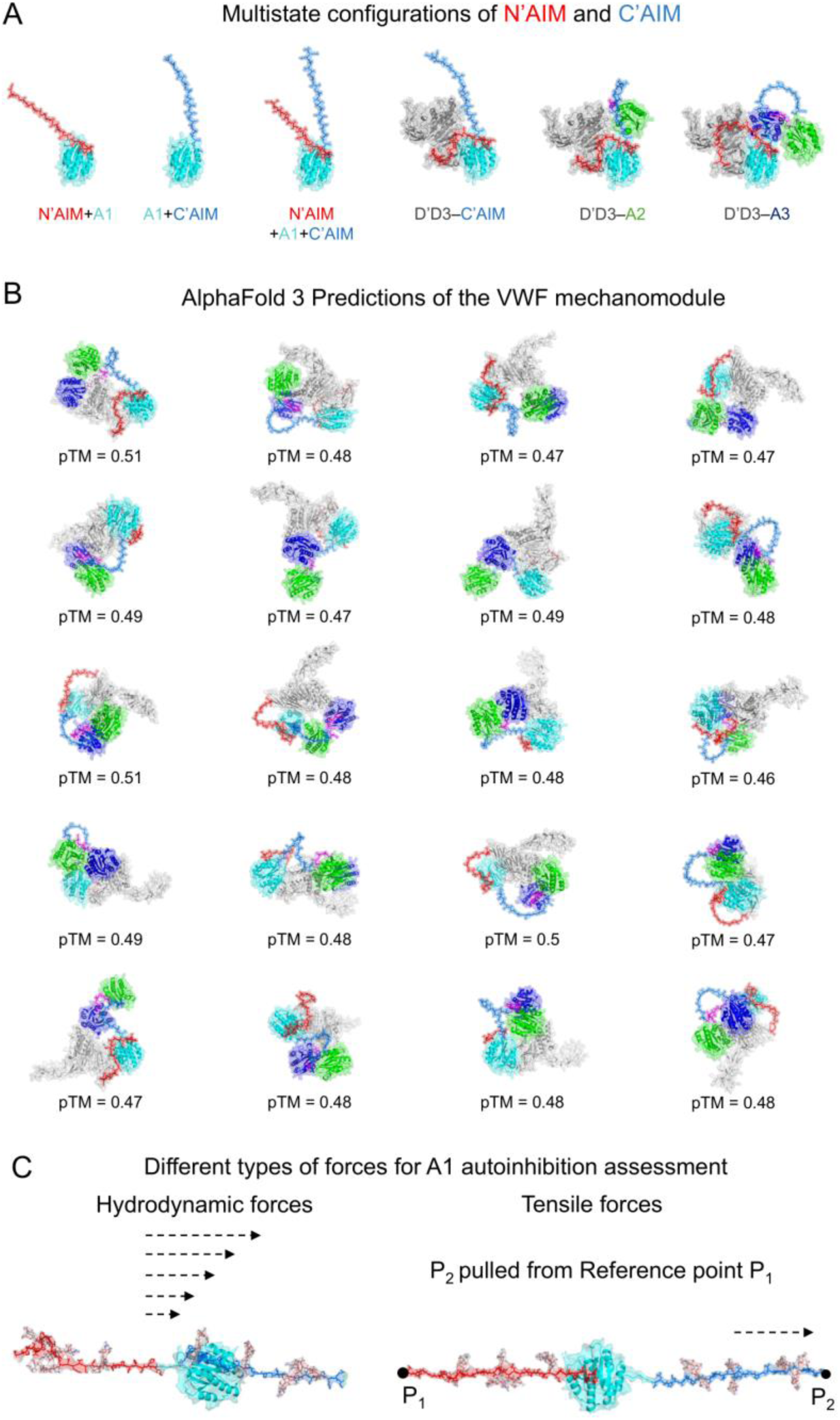
AlphaFold3 predictions of the VWF mechanomodule and structural configurations. (A) Diverse conformations of the N’AIM and C’AIM regions in the presence and absence of flanking neighboring domains. (B)Predicted structures of the VWF mechanomodule generated by AlphaFold3, accompanied by corresponding predicted Template Modeling (pTM) scores (n = 20 models). (C) Schematic overview highlighting the distinct types of mechanical forces used to study the alleviation of A1 autoinhibition and enable GPIbα binding: shear-like flow forces (left) and direct tensile forces (right).

**Fig. S2.**
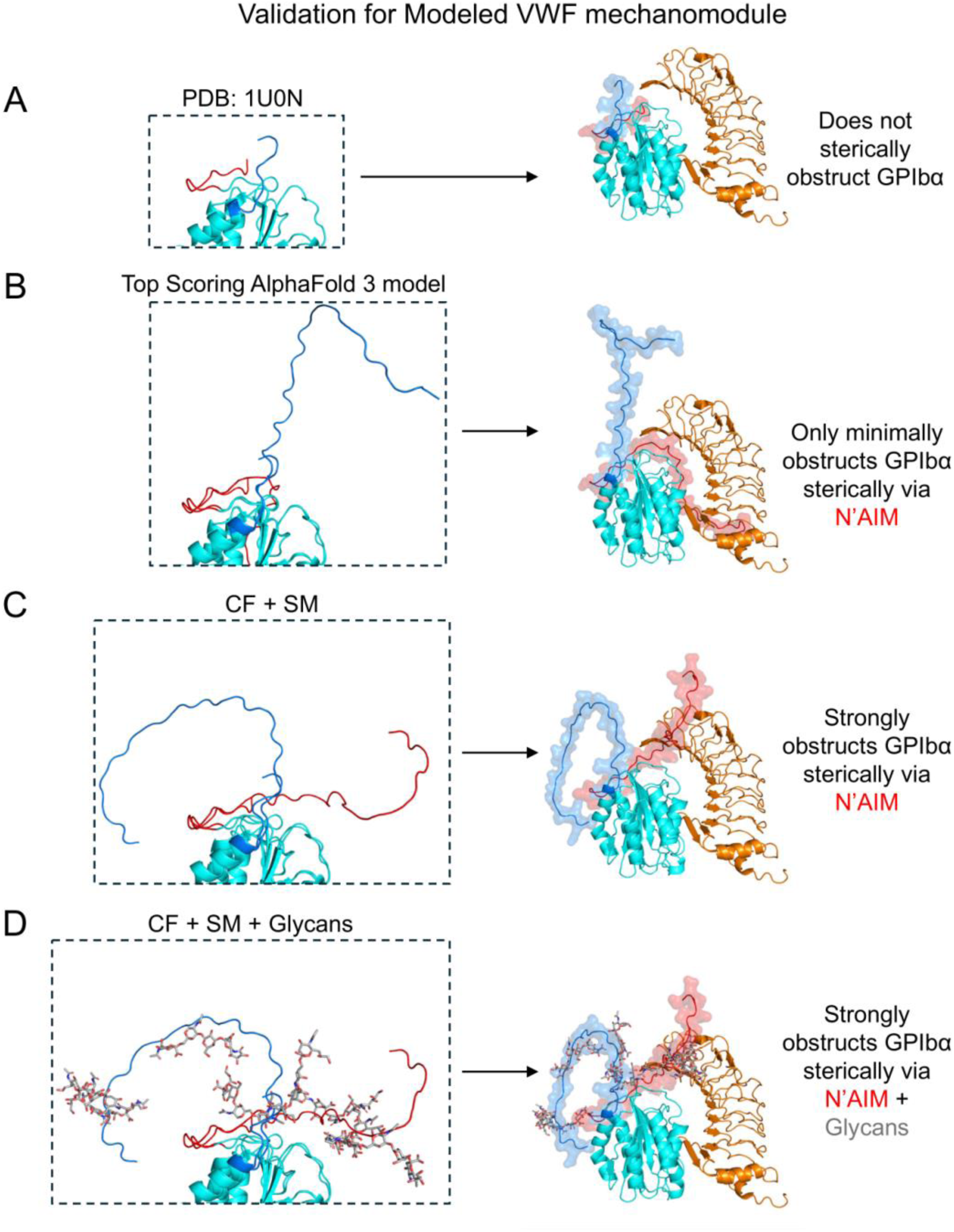
Validation of modeled VWF mechanomodule structures and their steric shielding of GPIbα. (A) Crystal structure of the A1 domain (PDB ID: 1U0N), containing the longest experimentally resolved N’AIM segment. (B) Top-scoring AlphaFold 3 prediction of the mechanomodule superimposed onto PDB 1SQ0 shows limited steric obstruction of the GPIbα binding interface by N’AIM. (C) (CF+SM) superimposed onto PDB 1SQ0 yields enhanced autoinhibitory conformation, with pronounced N’AIM-mediated obstruction of GPIbα access. (D) The glycosylated CF+SM model superimposed onto PDB 1SQ0 exhibits the highest degree of steric hindrance at the GPIbα-binding interface, highlighting the contribution of O-glycans to A1 autoinhibition

**Fig.S3.**
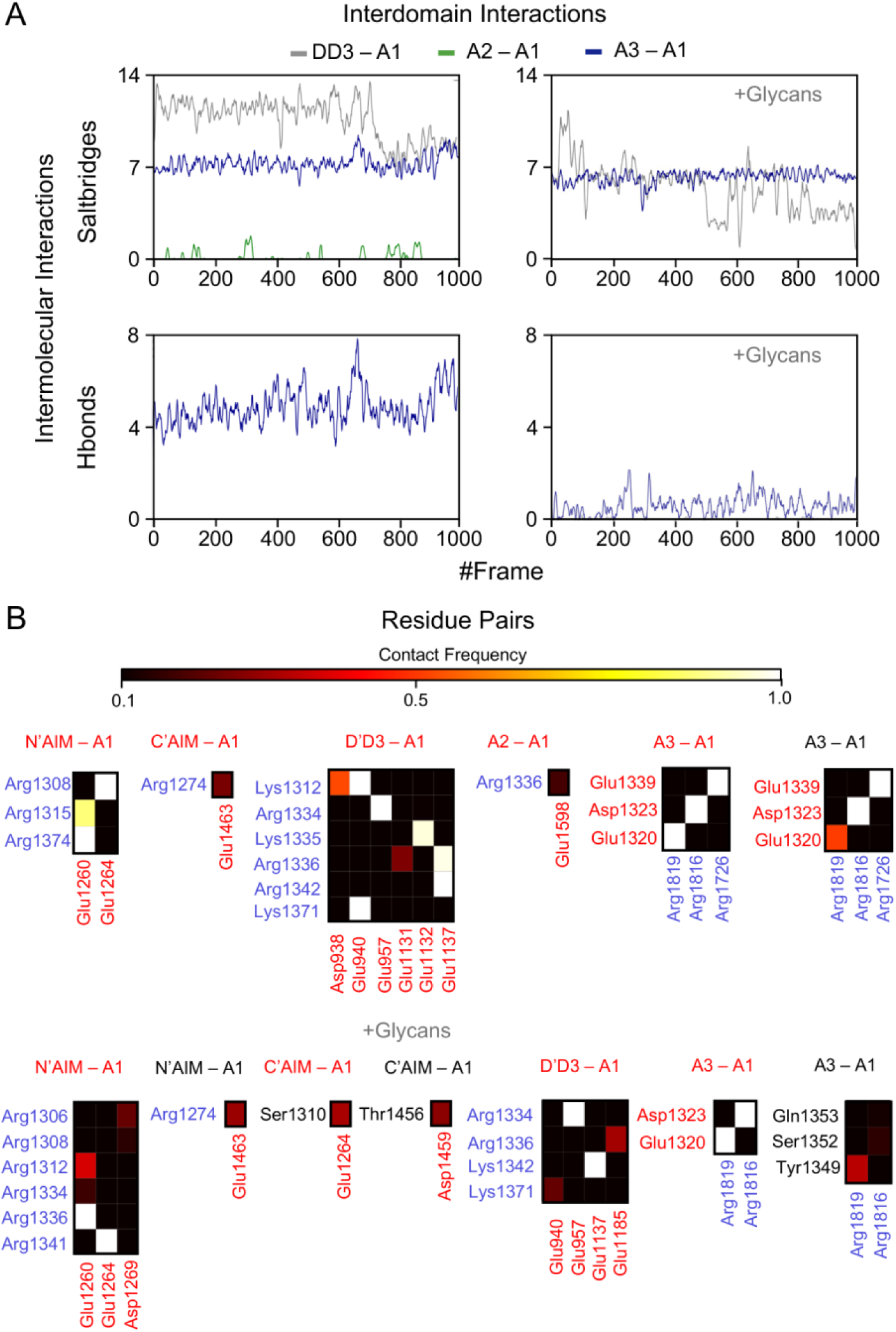
Intermolecular interactions of the VWF mechanomodule in the presence and absence of glycans. (A) Interdomain contacts between D’D3, A2, and A3 domains with the A1 domain observed during free molecular dynamics simulations. The presence of glycans introduces steric effects that modulate these interdomain interactions. (B) Contact frequency heatmap showing residue-residue interactions that persist in more than 10% of simulation frames. Low-frequency contacts are excluded. Brighter colors indicate higher contact frequencies, while darker colors represent transient or weaker interactions. (Positively charged residues are shown in blue, negatively charged residues in red, and all others in black; salt bridges are defined between oppositely charged residues)

**Fig.S4.**
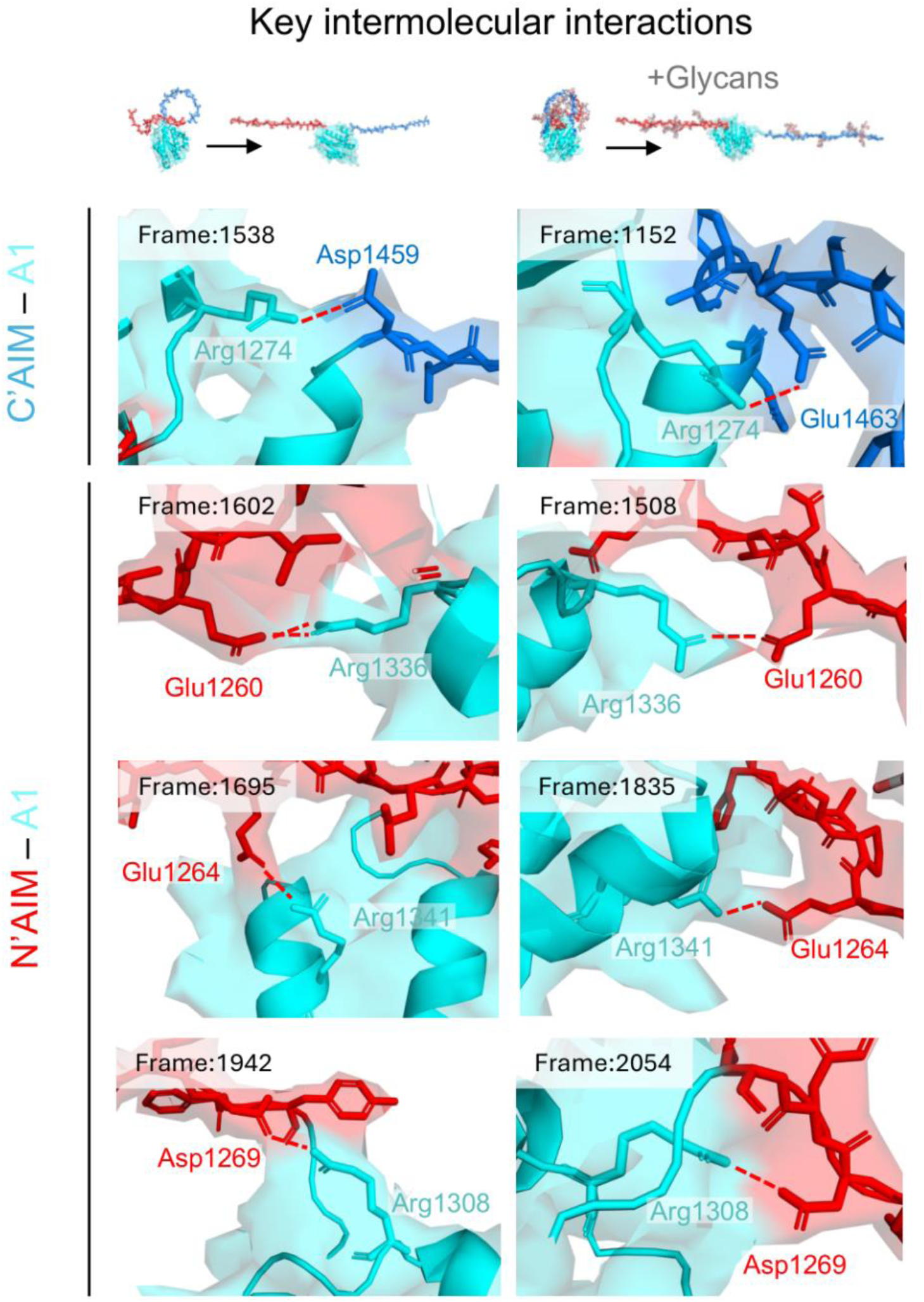
Trajectory snapshots of key residues mediating N′AIM and C′AIM engagement with the A1 domain from steered MD simulations. Snapshots illustrate how these interactions collectively influence AIM– A1 affinity in the presence (right) and absence (left) of O-linked glycans

**Fig.S5.**
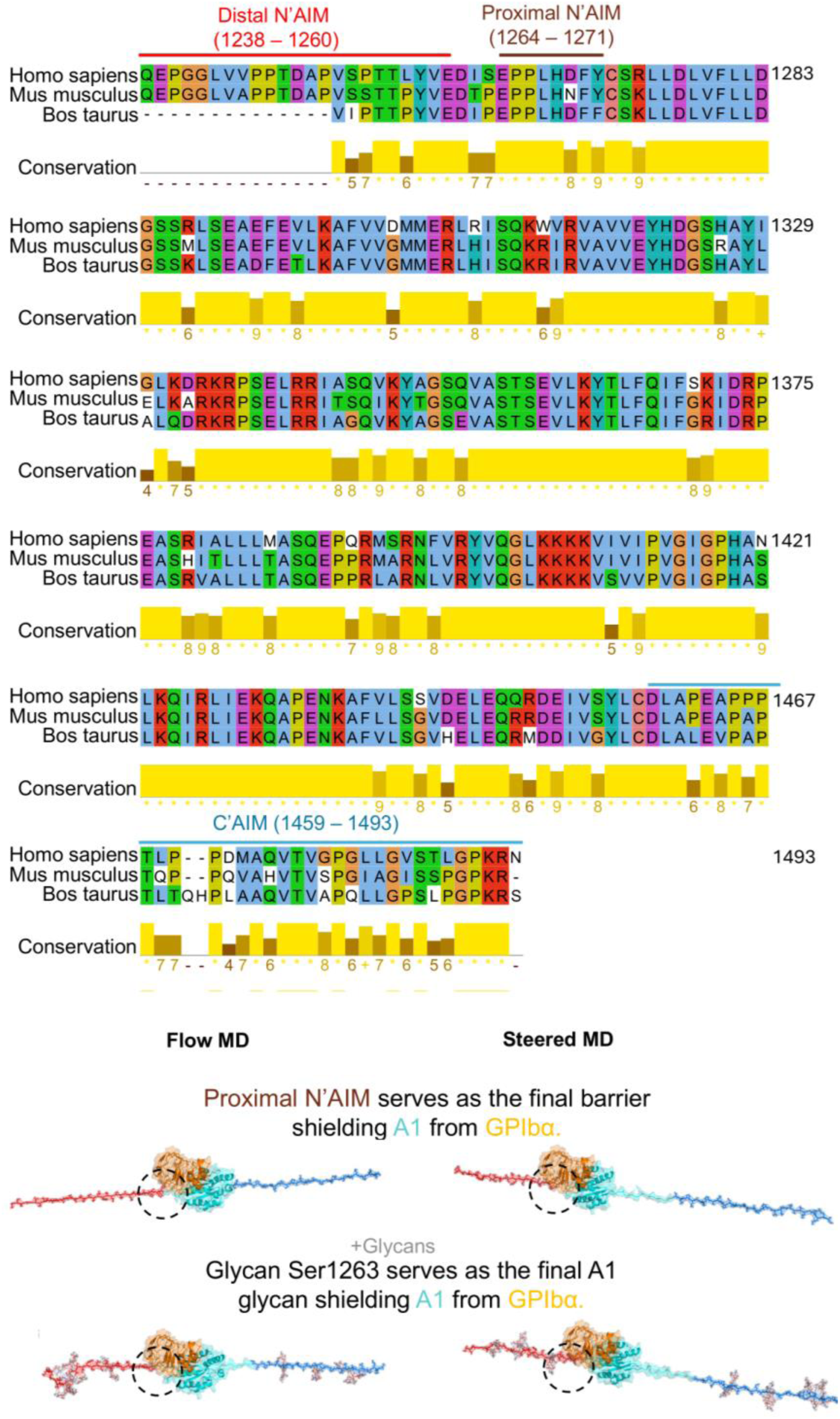
Conservation and structural analysis of the A1 domain across species. **(A)** Multiple sequence alignment of the N′AIM–A1–C′AIM regions of VWF comparing *Homo sapiens, Mus musculus,* and *Bos taurus*, revealing conserved and divergent residues relevant to autoinhibition and glycosylation. **(B)** Structural superimposition of PDB 1SQ0 onto A1 domains extracted from flow and steered MD trajectories, highlighting the final steric barriers to GPIbα access, specifically, the proximal N′AIM segment and the glycan at Ser1263

**Table S1.**
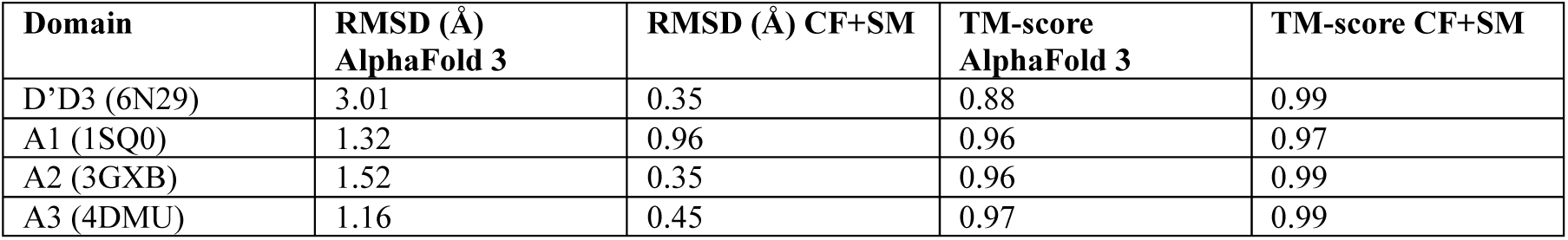
Structural similarity metrics (TM-score and RMSD) of VWF domains compared to crystal structures.

**Table S2.**
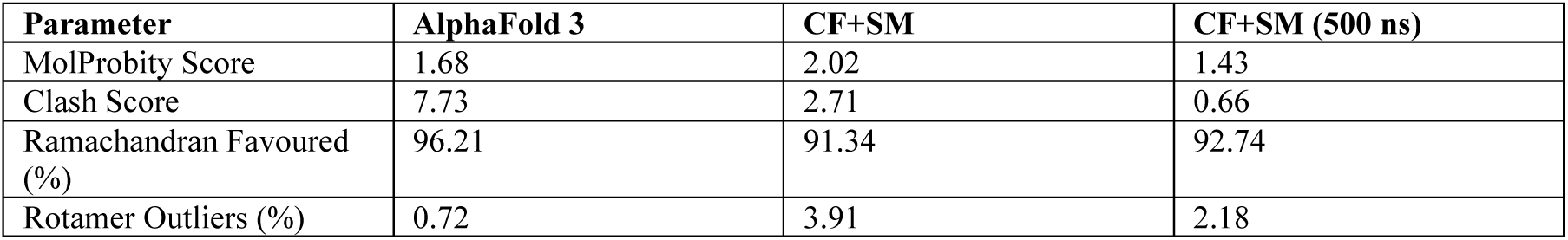
MolProbity structural validation scores of VWF mechanomodule models.

**Table S3.**
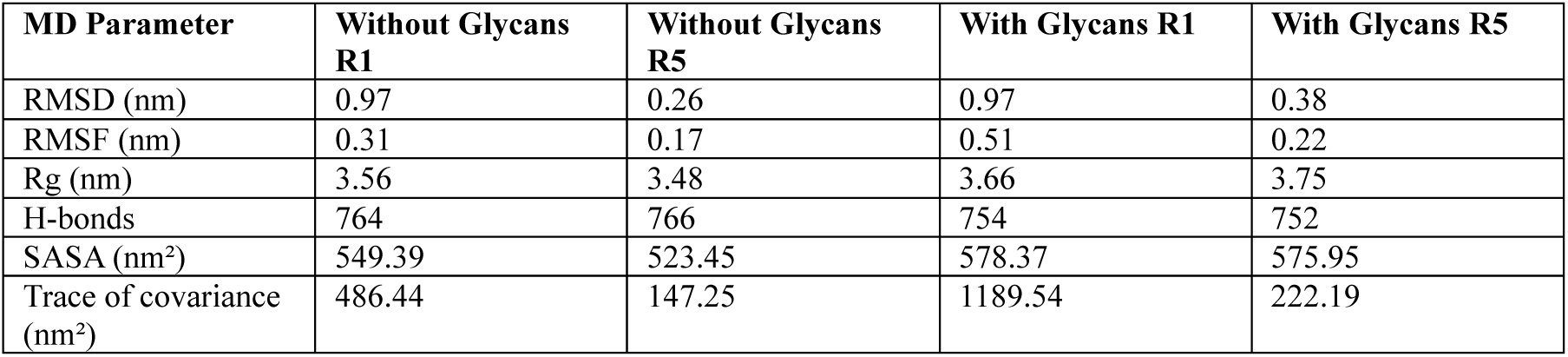
Average structural parameters from free MD simulations.

**Table S4.**
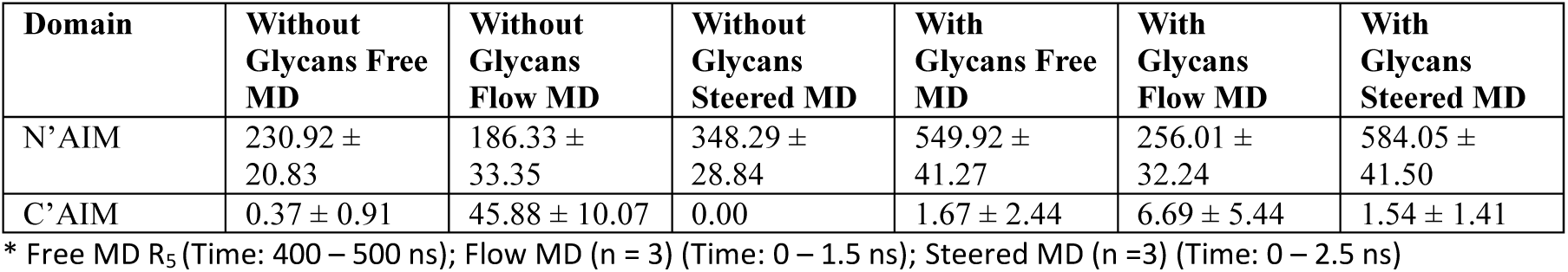
Average number of steric clashes between AIM and GPIbα (mean ± s.d.).

| Domain | Without Glycans Free MD | Without Glycans Flow MD | Without Glycans Steered MD | With Glycans Free MD | With Glycans Flow MD | With Glycans Steered MD |
| --- | --- | --- | --- | --- | --- | --- |
| N'AIM | 230.92 ± 20.83 | 186.33 ± 33.35 | 348.29 ± 28.84 | 549.92 ± 41.27 | 256.01 ± 32.24 | 584.05 ± 41.50 |
| C'AIM | 0.37 ± 0.91 | 45.88 ± 10.07 | 0.00 | 1.67 ± 2.44 | 6.69 ± 5.44 | 1.54 ± 1.41 |

\* Free MD R<sub>5</sub> (Time: 400 – 500 ns); Flow MD (n = 3) (Time: 0 – 1.5 ns); Steered MD (n =3) (Time: 0 – 2.5 ns)

**Table S5.** Average number of salt bridges (mean ± s.d.).

| Domain Pair | Without Glycans Free MD | Without Glycans Flow MD | Without Glycans Steered MD | With Glycans Free MD | With Glycans Flow MD | With Glycans Steered MD |
| --- | --- | --- | --- | --- | --- | --- |
| D'D3-A1 | 10.61 | 0.87 ± 0.23 | - | 5.47 | 1.21 ± 0.41 | - |
| N'AIM-A1 | 6.95 | 2.18 ± 0.36 | 4.14 ± 0.75 | 6.90 | 2.01 ± 0.52 | 3.11 ± 0.78 |
| C'AIM-A1 | 0.27 | 1.17 ± 0.64 | 0.05 ± 0.07 | 0.30 | 0.77 ± 0.61 | 0.06 ± 0.08 |
| N'AIM-C'AIM | 0.00 | 0.00 | - | 0.00 | 0.00 | - |
| A2-A1 | 0.18 | 0.00 | - | 0.00 | 0.00 | - |
| A3-A1 | 7.28 | 3.80 ± 0.47 | - | 6.31 | 1.98 ± 0.23 | - |

\* Free MD R<sub>5</sub> (Time: 400 – 500 ns); Flow MD (n = 3) (Time: 0 – 1.5 ns); Steered MD (n =3) (Time: 0 – 2.5 ns)

**Table S6.**
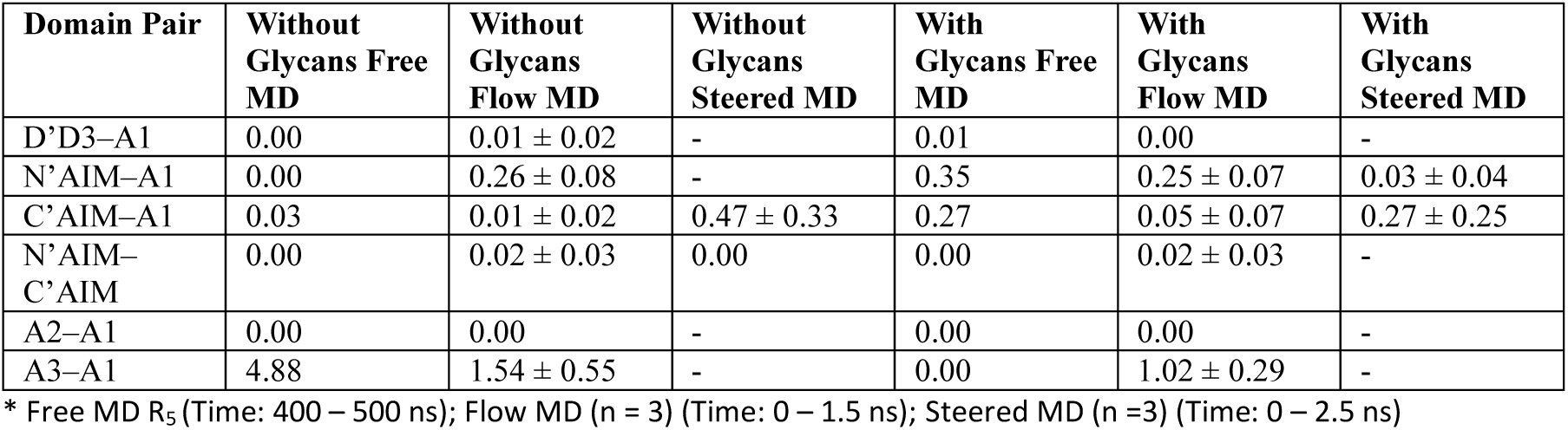
Average number of hydrogen bonds (mean ± s.d.).

| Domain Pair | Without Glycans Free MD | Without Glycans Flow MD | Without Glycans Steered MD | With Glycans Free MD | With Glycans Flow MD | With Glycans Steered MD |
| --- | --- | --- | --- | --- | --- | --- |
| D'D3-A1 | 0.00 | 0.01 ± 0.02 | - | 0.01 | 0.00 | - |
| N'AIM-A1 | 0.00 | 0.26 ± 0.08 | - | 0.35 | 0.25 ± 0.07 | 0.03 ± 0.04 |
| C'AIM-A1 | 0.03 | 0.01 ± 0.02 | 0.47 ± 0.33 | 0.27 | 0.05 ± 0.07 | 0.27 ± 0.25 |
| N'AIM-C'AIM | 0.00 | 0.02 ± 0.03 | 0.00 | 0.00 | 0.02 ± 0.03 | - |
| A2-A1 | 0.00 | 0.00 | - | 0.00 | 0.00 | - |
| A3-A1 | 4.88 | 1.54 ± 0.55 | - | 0.00 | 1.02 ± 0.29 | - |

\* Free MD R<sub>5</sub> (Time: 400 – 500 ns); Flow MD (n = 3) (Time: 0 – 1.5 ns); Steered MD (n =3) (Time: 0 – 2.5 ns)

## Notes

### Competing Interest Statement

The authors have declared no competing interest.

### Funding

- Digital Science Initiative 2024 Ignite Grant: M1IGNITE
- MRFF Cardiovascular Health Mission Grants
- MRFF Cardiovascular Health Mission Grants: MRF201616
- Snow Medical Research Foundation: 2022SF176
- Australian National Health and Medical Research Council: APP2003904
- National Heart Foundation of Australia (Heart Foundation): 106979
- National Computational Infrastructure (NCI): PTJ70
- Australian National Health and Medical Research Council: GNT2022247
- National Heart Foundation of Australia (Heart Foundation): 106879

