## Supplementary material for "Flow molecular dynamics simulations reveal mechanosensitive regulation of von Willebrand factor through glycan-modulated autoinhibitory modules": This PDF file includes Figures S1 to S5 and Tables S1 to S6

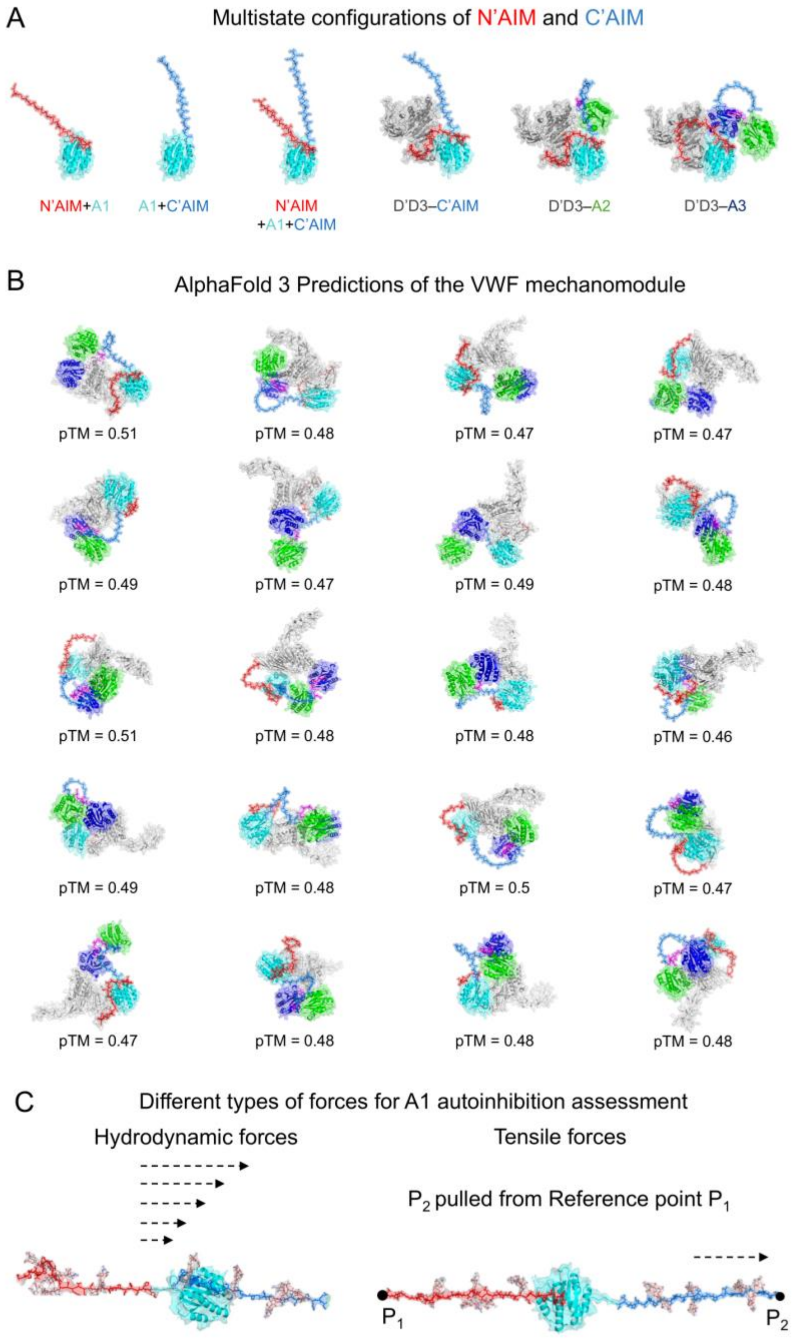


**Fig. S1** AlphaFold3 predictions of the VWF mechanomodule and structural configurations. (A) Diverse conformations of the N'AIM and C'AIM regions in the presence and absence of flanking neighboring domains. (B)Predicted structures of the VWF mechanomodule generated by AlphaFold3, accompanied by corresponding predicted Template Modeling (pTM) scores (n = 20 models). (C) Schematic overview highlighting the distinct types of mechanical forces used to study the alleviation of A1 autoinhibition and enable GPIbα binding: shear-like flow forces (left) and direct tensile forces (right).

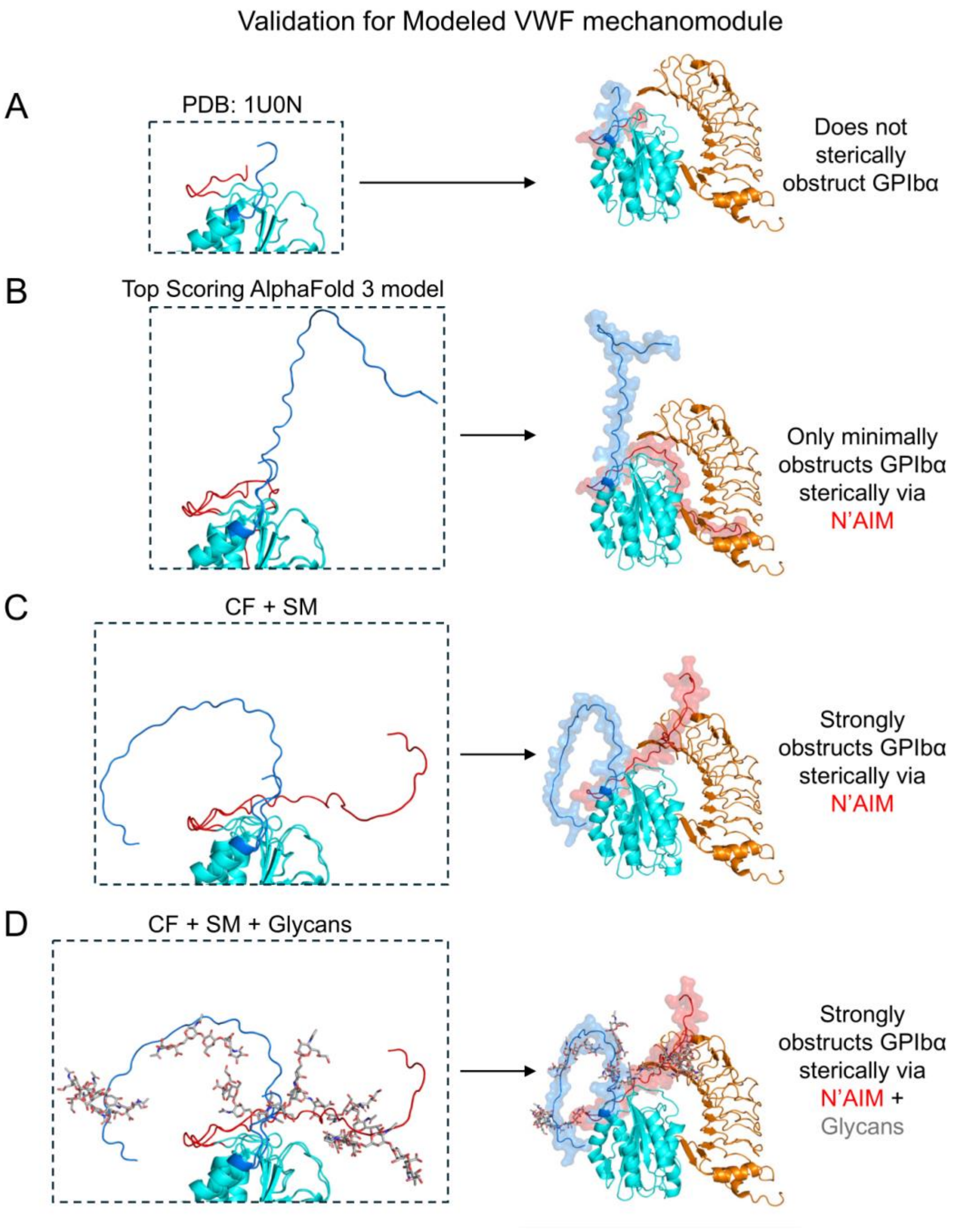


**Fig. S2** Validation of modeled VWF mechanomodule structures and their steric shielding of GPIbα. (A) Crystal structure of the A1 domain (PDB ID: 1U0N), containing the longest experimentally resolved N'AIM segment. (B) Top-scoring AlphaFold 3 prediction of the mechanomodule superimposed onto PDB 1SQ0 shows limited steric obstruction of the GPIbα binding interface by N'AIM. (C) (CF+SM) superimposed onto PDB 1SQ0 yields enhanced autoinhibitory conformation, with pronounced N'AIM-mediated obstruction of GPIbα access. (D) The glycosylated CF+SM model superimposed onto PDB 1SQ0 exhibits the highest degree of steric hindrance at the GPIbα-binding interface, highlighting the contribution of O-glycans to A1 autoinhibition

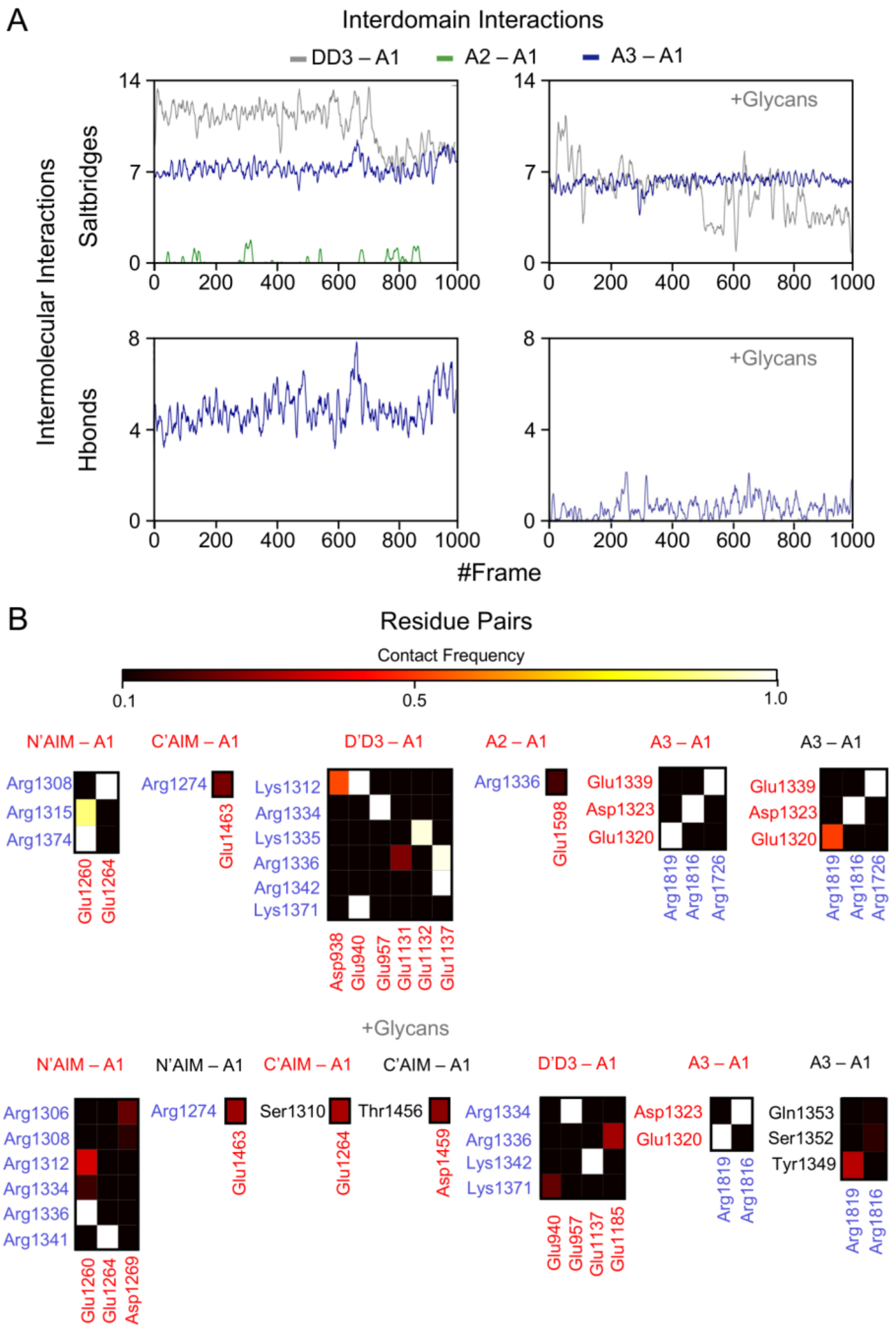


**Fig.S3** Intermolecular interactions of the VWF mechanomodule in the presence and absence of glycans. (A) Interdomain contacts between D'D3, A2, and A3 domains with the A1 domain observed during free molecular dynamics simulations. The presence of glycans introduces steric effects that modulate these interdomain interactions. (B) Contact frequency heatmap showing residue-residue interactions that persist in more than 10% of simulation frames. Low-frequency contacts are excluded. Brighter colors indicate higher contact frequencies, while darker colors represent transient or weaker interactions. (Positively charged residues are shown in blue, negatively charged residues in red, and all others in black; salt bridges are defined between oppositely charged residues)

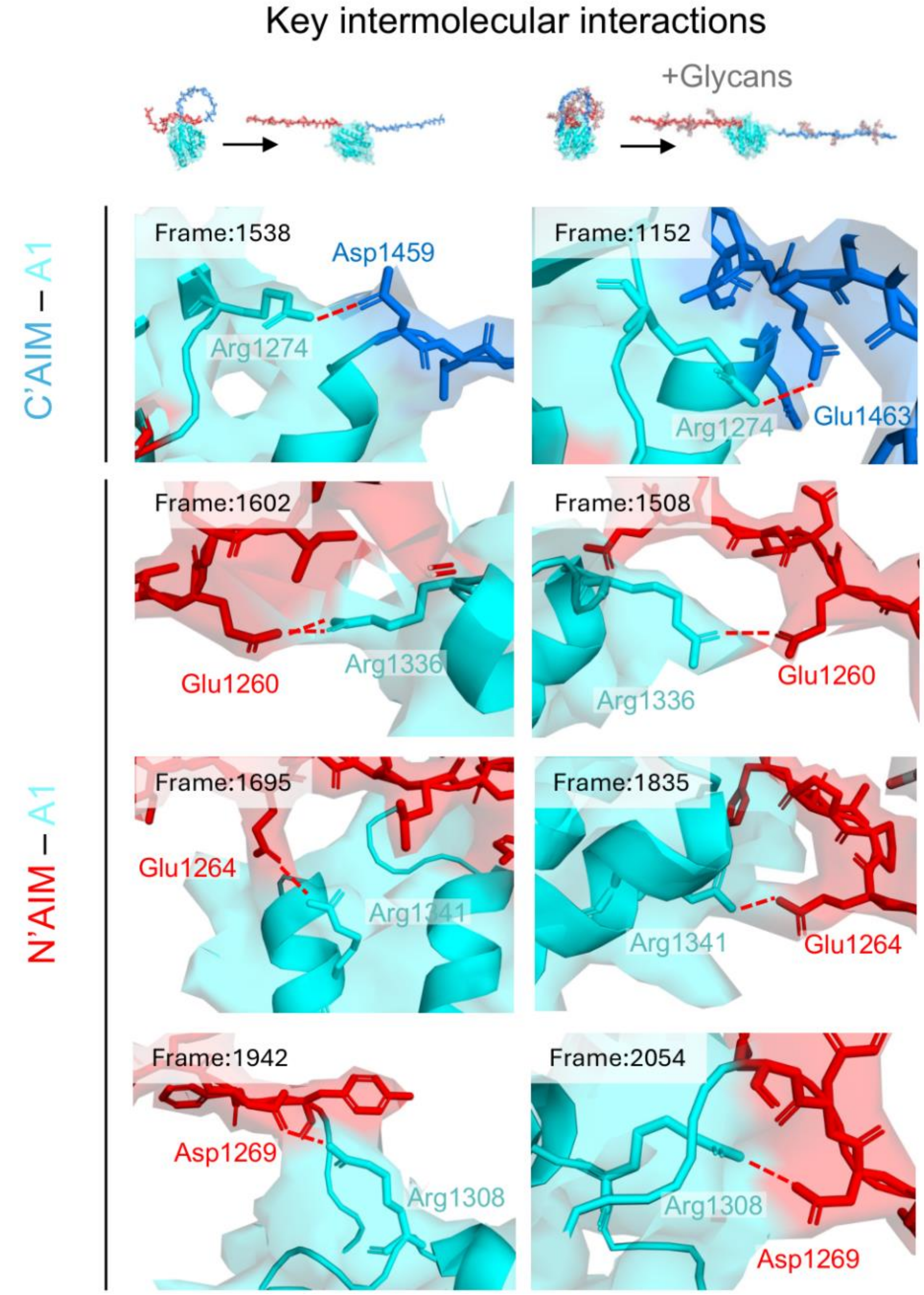


**Fig.S4 Trajectory snapshots of key residues mediating N′AIM and C′AIM engagement with the A1 domain from steered MD simulations.** Snapshots illustrate how these interactions collectively influence AIM– A1 affinity in the presence (right) and absence (left) of O-linked glycans

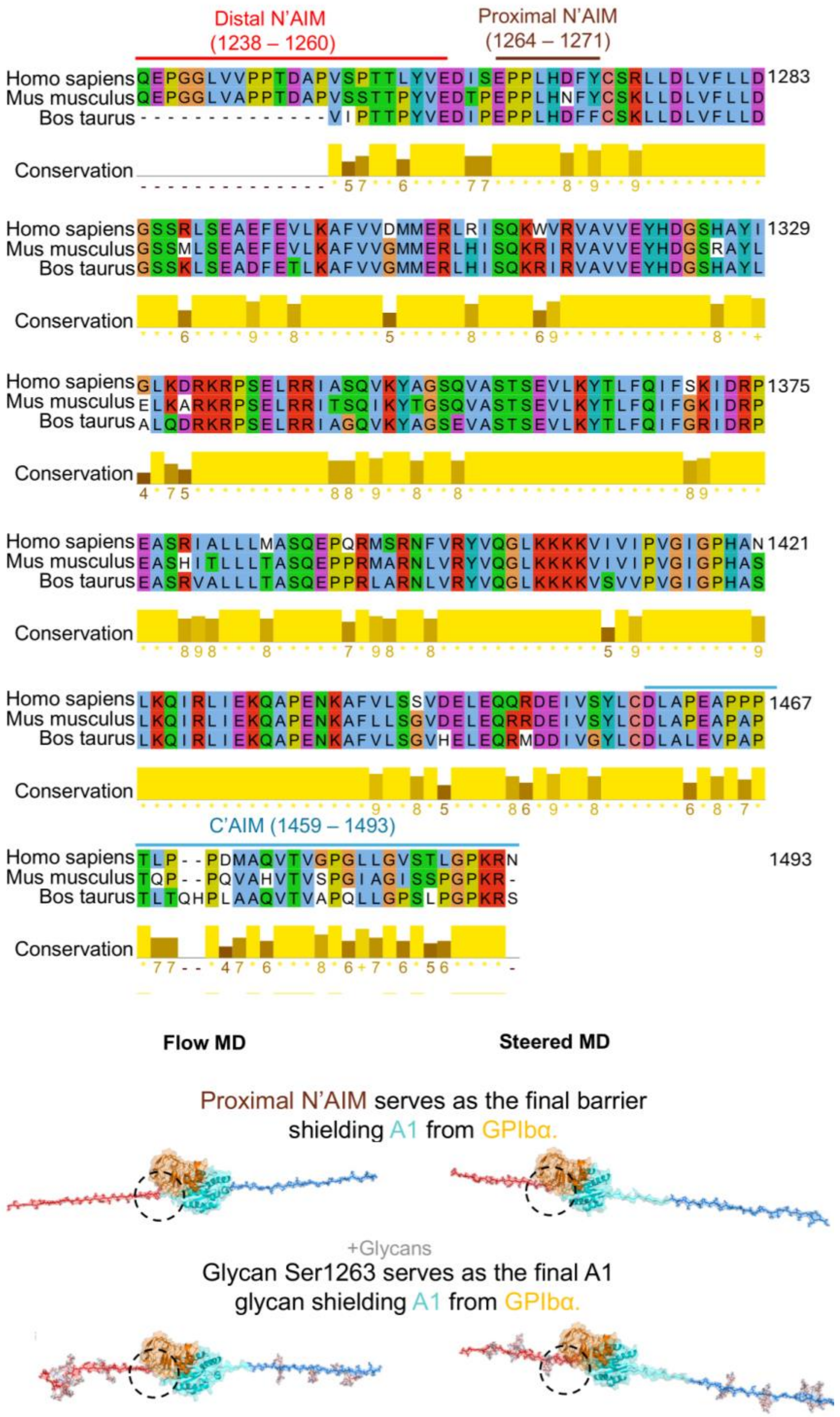


**Fig.S5 Conservation and structural analysis of the A1 domain across species. (A)** Multiple sequence alignment of the N′AIM–A1–C′AIM regions of VWF comparing *Homo sapiens, Mus musculus,* and *Bos taurus*, revealing conserved and divergent residues relevant to autoinhibition and glycosylation. **(B)** Structural superimposition of PDB 1SQ0 onto A1 domains extracted from flow and steered MD trajectories, highlighting the final steric barriers to GPIbα access, specifically, the proximal N′AIM segment and the glycan at Ser1263

### Tables

**Table S1.** Structural similarity metrics (TM-score and RMSD) of VWF domains compared to crystal structures.

| **Domain** | **RMSD (Å) AlphaFold 3** | **RMSD (Å) CF+SM** | **TM-score AlphaFold 3** | **TM-score CF+SM** |
|---|---|---|---|---|
| D'D3 (6N29) | 3.01 | 0.35 | 0.88 | 0.99 |
| A1 (1SQ0) | 1.32 | 0.96 | 0.96 | 0.97 |
| A2 (3GXB) | 1.52 | 0.35 | 0.96 | 0.99 |
| A3 (4DMU) | 1.16 | 0.45 | 0.97 | 0.99 |

**Table S2.** MolProbity structural validation scores of VWF mechanomodule models.

| Parameter | AlphaFold 3 | CF+SM | CF+SM (500 ns) |
|---|---|---|---|
| MolProbity Score | 1.68 | 2.02 | 1.43 |
| Clash Score | 7.73 | 2.71 | 0.66 |
| Ramachandran Favoured (%) | 96.21 | 91.34 | 92.74 |
| Rotamer Outliers (%) | 0.72 | 3.91 | 2.18 |

**Table S3.** Average structural parameters from free MD simulations.

| MD Parameter | Without Glycans R1 | Without Glycans R5 | With Glycans R1 | With Glycans R5 |
|---|---|---|---|---|
| RMSD (nm) | 0.97 | 0.26 | 0.97 | 0.38 |
| RMSF (nm) | 0.31 | 0.17 | 0.51 | 0.22 |
| Rg (nm) | 3.56 | 3.48 | 3.66 | 3.75 |
| H-bonds | 764 | 766 | 754 | 752 |
| SASA ($nm^2$) | 549.39 | 523.45 | 578.37 | 575.95 |
| Trace of covariance ($nm^2$) | 486.44 | 147.25 | 1189.54 | 222.19 |

**Table S4.** Average number of steric clashes between AIM and GPIbα (mean ± s.d.).

| Domain | Without Glycans Free MD | Without Glycans Flow MD | Without Glycans Steered MD | With Glycans Free MD | With Glycans Flow MD | With Glycans Steered MD |
|---|---|---|---|---|---|---|
| N'AIM | 230.92 ± 20.83 | 186.33 ± 33.35 | 348.29 ± 28.84 | 549.92 ± 41.27 | 256.01 ± 32.24 | 584.05 ± 41.50 |
| C'AIM | 0.37 ± 0.91 | 45.88 ± 10.07 | 0.00 | 1.67 ± 2.44 | 6.69 ± 5.44 | 1.54 ± 1.41 |

* Free MD $R_5$ (Time: 400 – 500 ns); Flow MD (n = 3) (Time: 0 – 1.5 ns); Steered MD (n =3) (Time: 0 – 2.5 ns)

**Table S5**. Average number of salt bridges (mean ± s.d.).

| Domain Pair | Without Glycans Free MD | Without Glycans Flow MD | Without Glycans Steered MD | With Glycans Free MD | With Glycans Flow MD | With Glycans Steered MD |
|---|---|---|---|---|---|---|
| D'D3–A1 | 10.61 | 0.87 ± 0.23 | - | 5.47 | 1.21 ± 0.41 | - |
| N'AIM–A1 | 6.95 | 2.18 ± 0.36 | 4.14 ± 0.75 | 6.90 | 2.01 ± 0.52 | 3.11 ± 0.78 |
| C'AIM–A1 | 0.27 | 1.17 ± 0.64 | 0.05 ± 0.07 | 0.30 | 0.77 ± 0.61 | 0.06 ± 0.08 |
| N'AIM–C'AIM | 0.00 | 0.00 | - | 0.00 | 0.00 | - |
| A2–A1 | 0.18 | 0.00 | - | 0.00 | 0.00 | - |
| A3–A1 | 7.28 | 3.80 ± 0.47 | - | 6.31 | 1.98 ± 0.23 | - |

* Free MD $R_5$ (Time: 400 – 500 ns); Flow MD (n = 3) (Time: 0 – 1.5 ns); Steered MD (n =3) (Time: 0 – 2.5 ns)

**Table S6.** Average number of hydrogen bonds (mean ± s.d.).

| Domain Pair | Without Glycans Free MD | Without Glycans Flow MD | Without Glycans Steered MD | With Glycans Free MD | With Glycans Flow MD | With Glycans Steered MD |
|---|---|---|---|---|---|---|
| D'D3–A1 | 0.00 | 0.01 ± 0.02 | - | 0.01 | 0.00 | - |
| N'AIM–A1 | 0.00 | 0.26 ± 0.08 | - | 0.35 | 0.25 ± 0.07 | 0.03 ± 0.04 |
| C'AIM–A1 | 0.03 | 0.01 ± 0.02 | 0.47 ± 0.33 | 0.27 | 0.05 ± 0.07 | 0.27 ± 0.25 |
| N'AIM–C'AIM | 0.00 | 0.02 ± 0.03 | 0.00 | 0.00 | 0.02 ± 0.03 | - |
| A2–A1 | 0.00 | 0.00 | - | 0.00 | 0.00 | - |
| A3–A1 | 4.88 | 1.54 ± 0.55 | - | 0.00 | 1.02 ± 0.29 | - |

* Free MD $R_5$ (Time: 400 – 500 ns); Flow MD (n = 3) (Time: 0 – 1.5 ns); Steered MD (n =3) (Time: 0 – 2.5 ns)
